# Uncertainty in Thermosensory Expectations Enhances an Illusion of Pain

**DOI:** 10.1101/2024.03.27.587070

**Authors:** Jesper Fischer Ehmsen, Niia Nikolova, Daniel Elmstrøm Christensen, Leah Banellis, Malthe Brændholt, Arthur S. Courtin, Camilla E. Kraenge, Alexandra G. Mitchell, Camila Sardeto Deolindo, Christian Steenkjær, Melina Vejlø, Christoph Mathys, Micah G. Allen, Francesca Fardo

## Abstract

The human brain has a remarkable ability to learn and update its beliefs about the world. Here, we investigate how thermosensory learning shapes our subjective experience of temperature and the misperception of pain in response to harmless thermal stimuli. Through computational modeling, we demonstrate that the brain uses a probabilistic predictive coding scheme to update beliefs about temperature changes based on their uncertainty. We find that these expectations directly modulate the perception of pain in the thermal grill illusion. Quantitative microstructural brain imaging revealed that the myeloarchitecture and iron content of the somatosensory cortex, the posterior insula and the amygdala reflect inter-individual variability in computational parameters related to learning and the degree to which uncertainty modulates illusory pain perception. Our findings offer a new framework to explain how the brain infers pain from innocuous thermal inputs. Our model has important implications for understanding the etiology of thermosensory symptoms in chronic pain conditions.

## Introduction

The ability to adapt to environmental changes and learn in the face of uncertainty is critical for generating precise and flexible responses to a wide range of stimuli. In the context of thermosensation and nociception, such adaptability allows us to effectively detect temperature shifts and avert potential tissue damage, even under conditions of incomplete or ambiguous information. This capability is not only essential for safeguarding our bodily integrity but also facilitates our interaction with an uncertain environment. Here, we report findings demonstrating that thermosensation relies on precision-weighted expectations, and that this extends to complex phenomena such as illusory pain, exemplified by the Thermal Grill Illusion (TGI).

Current knowledge of the thermosensory and thermo-nociceptive systems predominantly revolves around peripheral sensory mechanisms that transduce innocuous and noxious thermal stimuli into neural signals. This includes landmark discoveries like the TRPV1 and TRPM8 receptors [1–3]. While these bottom-up mechanisms have been extensively studied, less attention has been devoted to how they integrate with top-down expectations to form our subjective experiences of temperature and pain. Indeed, perception in these domains is not solely the output of isolated afferent channels but is heavily influenced by prior beliefs and expectations [4–7]. In this context, the TGI presents a striking case in which the simultaneous presentation of innocuous warm and cold stimuli can evoke illusory burning sensations [12]. The illusion’s occurrence, which cannot be solely attributed to the physical characteristics of the stimuli, suggests that top-down expectations may shape the veridical perception of temperature and pain.

Associative learning plays a fundamental role in the perception of pain and its modulation by expectation, enabling the development of adaptive behaviors that protect us from potential harm. Significant progress has been made in understanding these processes through the computational neuroscience of predictive coding [13–22] and reinforcement learning [18,23]. For instance, it has been shown that participants learn about painful stimuli in a manner that is consistent with Bayesian principles [23,24], and pain-prediction errors have been mapped to key brain areas involved in pain-related processing, including the insula and brainstem [15,25,26]. A key contribution of this work was the recognition that expectation-related modulation of pain, such as nocebo and placebo effects [27–33], are grounded in the weighting of pain prediction errors by their uncertainty or inverse precision [13,18,34–36]. To date, it is unknown if these principles similarly explain innocuous thermosensory perception and illusions of pain. An intriguing possibility is that the TGI may stem from thermosensory predictive coding, where increased uncertainty about upcoming stimulus temperatures give rise to the perception of pain.

In this study, we apply computational methods to reveal how both innocuous thermosensation and the TGI are shaped by uncertainty in a large cohort of healthy participants. We further utilized high-resolution quantitative MRI to identify how inter-individual variations in brain microstructure are associated with computational fingerprints of thermosensory learning. To this aim, we conducted an experiment in 267 participants who completed a probabilistic thermosensory learning (PTL) task, in which we strategically embedded simultaneous cold and warm stimuli to induce the TGI within the learning sequence. This experimental approach offers a comprehensive analysis of the role of expectations in thermosensory learning and provides a unique opportunity to test the hypothesis that the uncertainty of thermal expectations plays a crucial role in the perception of illusory pain. Our results provide a compelling example of how precision-weighted expectations can lead to the misinterpretation of non-nociceptive stimuli as painful, offering potential new insights into symptoms of neuropathic and nociplastic pain conditions [37,38].

## Results

To quantify the relationship between learned expectations and thermosensation, we tested a novel probabilistic thermal learning task (PTL, Fig 1A) in 267 healthy individuals. The PTL integrates key features of reversal learning tasks in other sensory domains [39–42], in which participants must dynamically update sensory predictions in response to varying uncertainty. In each PTL trial, participants heard auditory cues consisting of high or low tones that predicted whether the forthcoming stimulus would be cold or warm. Critically, these cue-stimulus pairings shifted unpredictably over time, requiring participants to continuously relearn their associative mappings. Cue-stimulus associations varied according to blocks of longer, more stable periods in which reversals were less likely, and shorter, more variable periods in which transitions occurred more frequently (Fig 1B). Innocuous cool and warm trials were pseudorandomly interspersed with ambiguous stimuli. These ambiguous stimuli were the simultaneous presentation of the same objective temperatures as those used in the innocuous cold and warm trials, in an alternated spatial configuration. This method of stimulus presentation is known to elicit burning pain sensations referred to as the Thermal Grill Illusion (TGI)[8,10]. On each trial, participants made a binary prediction response, indicating whether they expected an upcoming cold or warm stimulus. In a subset of trials, they subsequently provided visual analog scale (VAS) ratings reflecting their perceived levels of cold, warm, and burning sensations.

**Fig 1.**
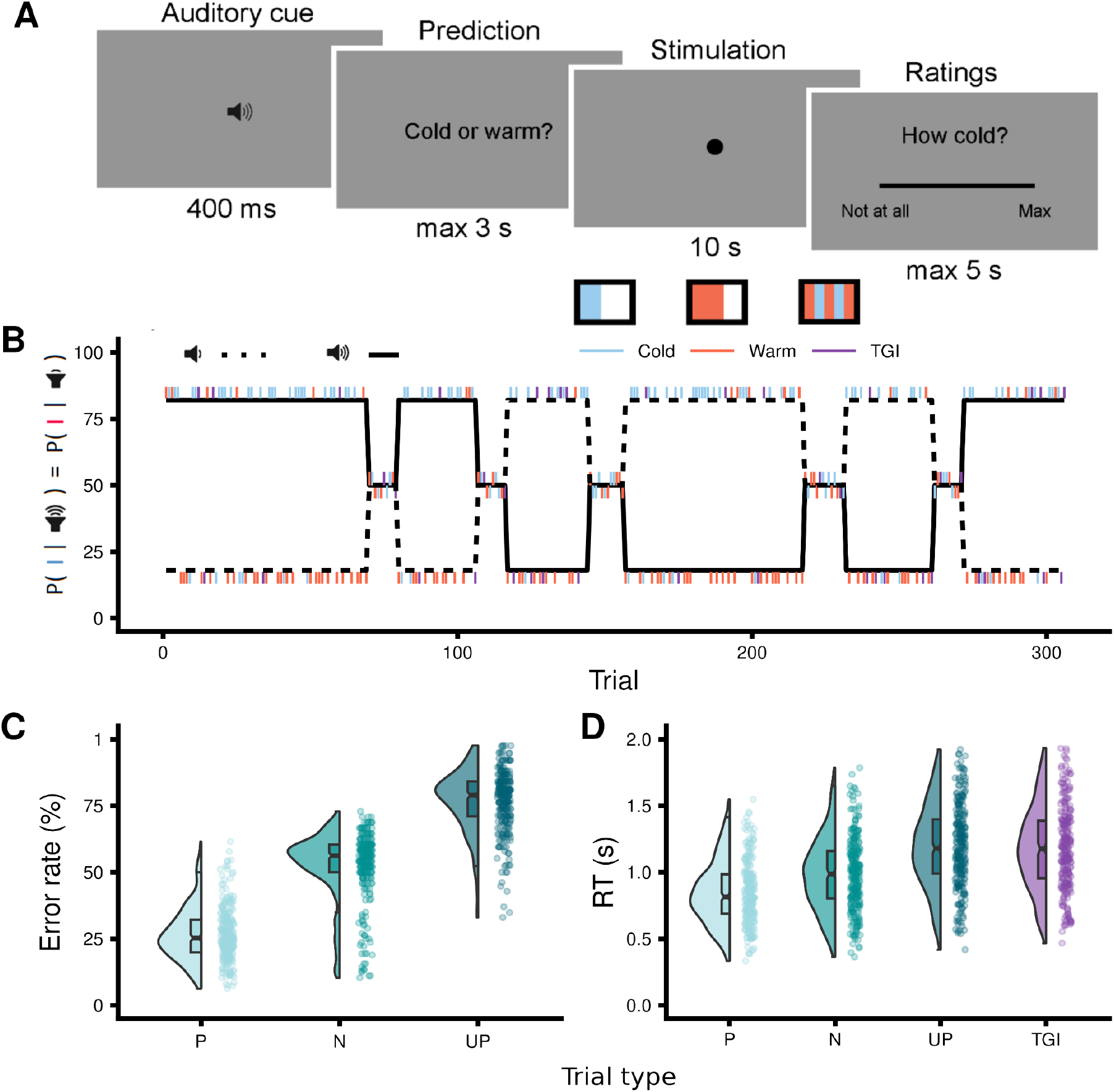
Thermosensory learning: experimental design and behavioral measures. **A.** Trial structure depicting the sequence of events within each trial: auditory cue presentation, prediction of the forthcoming stimulation quality as either cold or warm, delivery of the thermal stimulation (cold, warm or TGI) and VAS ratings of cold, warm and burning sensations. All three ratings were completed for a given stimulus. **B.** Time-course of cue-stimulus contingencies throughout the experiment, varying across three levels of cue-stimulus association probabilities set at 82%, 50% and 18%. **C.** Comparison of error rates for participants’ predictions of the forthcoming stimulation quality across predicted (P), neutral (N) and unpredicted (UP) innocuous thermosensory trials. **D.** Comparison of response times in the trial following predicted (P), neutral (N) and unpredicted (UP) thermosensory stimuli, as well as TGI stimuli, demonstrating post-prediction error slowing.

### Behavior

#### Error rates and response times are modulated by thermosensory learning

To evaluate participants’ learning of cue-stimulus associations, we analyzed error rates for predicted, neutral and unpredicted innocuous thermosensory stimuli (Fig 1C and Supplementary Table 1A). Predicted and unpredicted stimuli were defined based on the participants’ trial-by-trial predictions (i.e. whether they predicted a cold or a warm stimulus) in blocks where the nominal probability of a specific cue-stimulus association was 82% and 18%, respectively. Neutral trials referred to non-predictive blocks where a cue predicted a particular stimulus with a 50% probability. This analysis confirmed that the probability of cue-stimulus association robustly modulated expectations such that participants’ prediction accuracy was highest for predicted trials compared to both neutral (β = −1.34, 95% CI = [−1.38; −1.29], p < .0001) and unpredicted trials (β = −2.26, 95% CI = [−2.31; −2.21], p < .0001).

As further evidence of successful learning, we observed post-prediction error slowing, indicated by reduced response times on trials following association violations (Fig 1D and Supplementary Table 1B). Our findings showed that response times were increasingly slowed following neutral (β = 0.15, 95% CI = [0.13; 0.16], p < .0001) unpredicted (β = 0.31, 95% CI = [0.3; 0.32], p < .0001), and TGI stimuli (β = 0.34, 95% CI = [0.32; 0.35], p < .0001), compared to predicted stimuli. Together, these results serve as a model-free positive control, confirming that participants effectively learned and incorporated cue-stimulus relationships into their thermosensory predictions.

#### Stimulus-Specific Effects on Thermosensory and Burning Ratings

To evaluate the effectiveness of cold, warm and TGI stimuli, we predicted subjective ratings using linear mixed effects models incorporating a zero-one inflated beta regression approach (Supplementary Note). The TGI is characterized by an enhanced perception of heat and the elicitation of burning sensations when innocuous cold and warm stimuli are combined, which does not occur when these stimuli are applied individually (Fig 2, Supplementary Tables 2A-C). In line with heat enhancement, TGI stimuli were rated as significantly less cold than innocuous cold stimuli (β = −0.39, 95% CI = [−0.41; −0.37], p < .0001), but warmer than innocuous warm alone (β = 0.18, 95% CI = [0.17; 0.2], p < .0001). Further, in line with the elicitation of illusory pain, the concurrent application of cold and warm stimuli during TGI produced significantly greater burning sensations than when either cold (β = −0.45, 95% CI = [−0.48; −0.43], p < .0001), or warm (β = −0.65, 95% CI = [−0.68; −0.63], p < .0001) were applied individually. Taken together, these findings confirm that innocuous thermosensory stimuli were perceived in a veridical manner, and the TGI manipulation effectively induced illusory heat and burning sensations.

**Fig 2.**
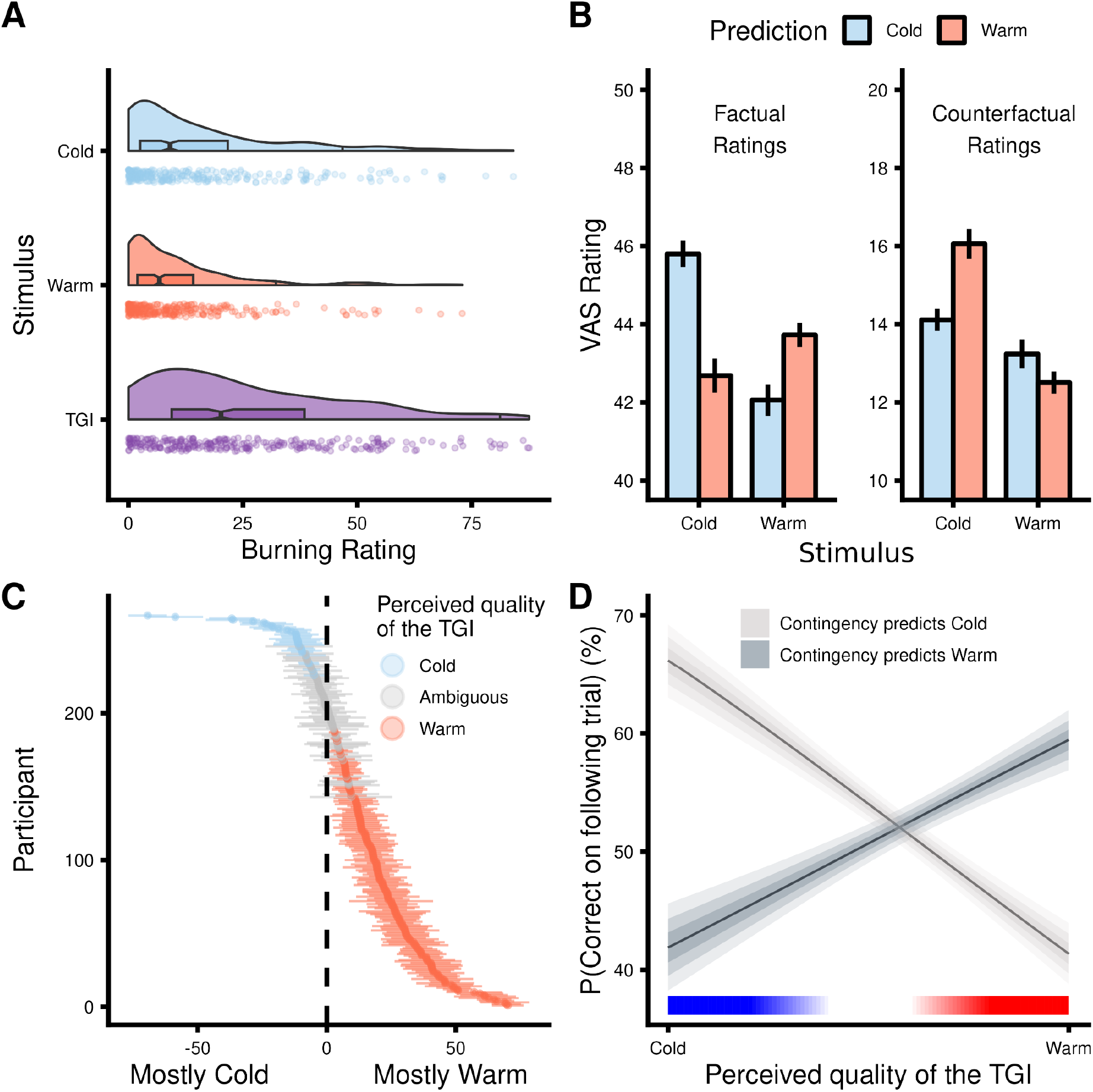
Thermosensory ratings and TGI perception. **A.** VAS burning ratings for cold, warm and TGI stimuli, illustrating a key feature of the TGI as an illusion of pain. **B.** Effects of participants’ expectations on VAS ratings for innocuous cold and warm stimuli, showing that expectations modulated these sensations (mean ± 2 SEM). **C.** Within-subject consistency of TGI perception as mostly cold (blue), ambiguous (gray) or mostly warm (red). Thermal ambiguity in this context signified that participants perceived the TGI trials as equally warm and cold. The y-axis depicts each individual participant, while the x-axis represents the ratio of perceived coldness to warmth for TGI stimuli (mean ± 2 SEM). **D.** The relationship between perceived TGI quality and learning (i.e., error rates in the trials that followed TGI stimulation), demonstrating that TGI trials reinforced cue-stimulus contingencies based on the perceived TGI quality.

#### Innocuous thermosensation is shaped by expectations

To investigate the impact of learned expectations on innocuous thermosensory experiences, we analyzed participants’ reported levels of both cold and warm sensations for predicted and unpredicted stimuli (Fig 2B and Supplementary Table 2D). For each stimulus, participants provided ratings for factual (e.g., coldness of a cold stimulus) and counterfactual qualities (e.g., warmth of a cold stimulus) of their sensations. We found a robust three-way interaction between the stimulation quality, the participants’ prediction on a trial by trial basis, and the rating type (β = 0.24, 95% CI = [0.17; 0.32], p < .0001). Considering the factual ratings, predicted cold stimuli were rated as colder than unpredicted cold stimuli (β = −0.09, 95% CI = [−0.12; −0.07], p < .0001), and predicted warm stimuli were rated as warmer than unpredicted warm stimuli (β = −0.05, 95% CI = [−0.07; −0.03], p < .0001). Conversely, when assessing the counterfactual quality, predicted cold stimuli were rated as less warm compared to the unpredicted stimuli (β = 0.06, 95% CI = [0.01; 0.11], p < .05), while predicted and unpredicted warm stimuli were rated as similarly cold (β = 0.04, 95% CI = [−0.01; 0.09], p = 0.13). Overall, these results highlight that participants’ thermosensory expectations significantly influenced the perceived intensity of innocuous stimuli.

#### Response times and error rates reflect perceived TGI quality

We hypothesized that the ambiguous nature of TGI trials would either reinforce or counter cue-stimulus associations, depending on the participants’ perception of TGI as primarily warm or cold. For instance, if a participant associates a high tone with a high probability of experiencing a cold stimulus, and perceives a TGI stimulus as predominantly warm, they might incorrectly infer a reversal has occurred after hearing a high tone and receiving a TGI stimulus, leading to an erroneous prediction in the subsequent trial. Conversely, if a participant perceives the TGI as predominantly cold, the participant’s correct association would be reinforced, leading to increased likelihood of an accurate prediction in the subsequent trial. To evaluate this hypothesis, we assessed each participant’s perceived TGI quality by computing the ratio of perceived coldness to warmth. In general, participants displayed high self-consistency in evaluating their perception of TGI stimuli as predominantly cold or warm (Fig 2C).

Our model confirmed our hypothesis, demonstrating that error rates were significantly influenced by the interaction between cue-stimulus association and perceived TGI quality (β = −1.73, 95% CI = [−2.04; −1.42], p < .0001, Fig 2 and Supplementary Table 2E). Specifically, participants were more likely to respond correctly on the subsequent trial when the contingency and the perceived TGI quality matched (i.e., predicting cold and perceiving TGI as predominantly cold) (β = 1.02, 95% CI = [0.81; 1.24], p < .0001). Conversely, participants were more likely to make incorrect responses in the following trial when the contingency and the perceived TGI quality diverged (e.g., predicting warm and perceiving TGI as predominantly cold) (β = −0.71, 95% CI = [−0.94; −0.48], p < .0001). Collectively, in a contingency block that predicted a cold outcome, the odds of making a correct prediction changed substantially depending on whether the TGI was rated as mostly cold vs. warm - a difference amounting to a 18.36% change in the probability of a correct answer [22.13, 14.08%]. Complementary effects were observed for response times after TGI stimulation (see Supplementary Results and Supplementary Table 2F). In summary, these findings reveal that TGI trials play a crucial role in reinforcing cue-stimulus associations by effectively shaping participants’ thermosensory predictions based on their perceived quality.

### Computational modeling

#### A 2-level Hierarchical Gaussian Filter model best explained thermosensory learning

We employed the Hierarchical Gaussian Filter (HGF) [43,44] to analyze learning trajectories across two hierarchical levels of belief (Fig 3). We estimated mean and uncertainty values for beliefs about an upcoming stimulus given a cue (i.e., predictions) and beliefs regarding the strength of cue-outcome associations (i.e., estimations). At the first level (*x*_1_), prediction uncertainty, pertains to uncertainty about immediate outcomes. A low prediction uncertainty indicates high confidence in predicting the forthcoming stimulus based on the given cue, while high prediction uncertainty suggests that the participant has not formed a definite prediction of which outcome is most likely. At the second level (*x*_2_), estimation uncertainty, quantifies the uncertainty surrounding the reliability of cue-outcome relationships. This level of uncertainty influences the rate at which beliefs about cue-outcome associations are updated. Low estimation uncertainty signifies a strong belief in the consistency of the cue-outcome association, requiring considerable contrary evidence for a belief update. In contrast, high estimation uncertainty means that beliefs regarding the cue-outcome relationship are more malleable and can be adjusted more readily upon encountering disconfirming evidence. Prediction and estimation uncertainty are structured hierarchically, meaning that beliefs at one level are dependent on, or informed by, the beliefs at the upper level (Fig 3).

**Fig 3.**
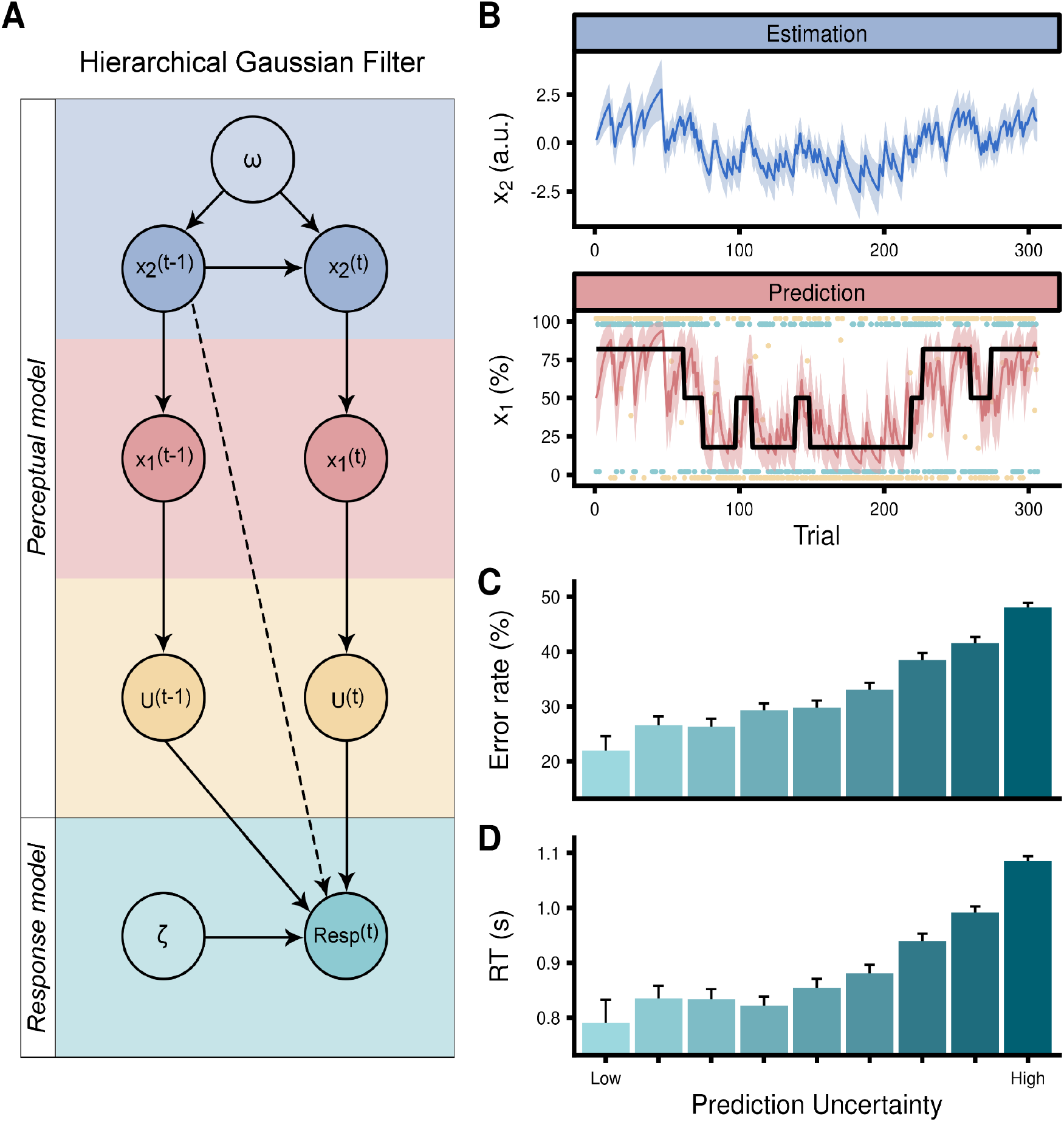
Computational modeling of thermosensation and illusory pain. **A.** Illustration of the Hierarchical Gaussian Filter, and its constituent perceptual and response models. Within the perceptual model, two hierarchical levels of trajectories with uncertainties are defined: prediction (*x*_1_) and estimation (*x*_2_). The first level takes the form of a Bernoulli distribution, while the second level evolves in time as a Gaussian random walk with step-size corresponding to the omega (ω) parameter. The response model converts the continually updated perceptual belief to a probability of answering through the inverse decision temperature zeta (ζ) through a logistic sigmoid transformation. U are observed values representing the cue-stimulus association mappings, dashed line depicts mediation through model inversion. **B.** Example of a single participants’ prediction and estimation trajectories together with their respective uncertainties. When considering the prediction trajectory, the thick black line represents the actual contingency probabilities. The trial-by-trial participant’s responses (i.e., predictions) are depicted by green points, where the value of one corresponds to the prediction of a cold stimulus and the value of zero corresponds to a prediction of a warm stimulus. The contingency space is represented by yellow dots, where zero values represent low tone-cold and high tone-warm associations and one values represent low tone-warm and high tone-cold associations. Intermediate values represent trials in which the stimulus was simultaneously cold and warm (i.e., TGI). Prediction uncertainty strongly modulated both **C.** error rates and **D.** response times needed to provide a prediction about the upcoming stimulus, validating the response model. Prediction uncertainty is presented here as discretized into nine bins.

To assess the best-fitting model, while accounting for parameter complexity, we compared the 2-level HGF with other well-known well-known reinforcement learning models, such as Rescorla-Wagner and Sutton K1 using Bayesian model selection [45]. We found that the 2-level HGF outperformed these models. To validate the robustness of our fitted models, we conducted both parameter and model recovery for the models under consideration (see Supplementary Figures S1-S5). Overall, model comparison and cross-validation demonstrated that thermosensory learning is best captured by Bayesian precision-weighted mechanisms that integrate both prediction and estimation uncertainty.

#### Modulation of behavior and perception by uncertainty

The impact of uncertainty on behavior and subjective experience was assessed using hierarchical regression analyses. At the lower level, involving prediction uncertainty, precise beliefs notably diminished error rates (β = −4.88, 95% CI = [−5.13; −4.63], p < .0001, Fig 3C and Supplementary Table 3A) and response times (β = 1.76, 95% CI = [1.69; 1.82], p < .0001, Fig 3d and Supplementary Table 3B) when participants predicted the quality of a forthcoming stimulus. This effect was also reflected in heightened VAS ratings for the thermosensory quality consistent with participants’ expectations (β = −0.19, 95% CI = [−0.28; −0.09], p < .0001). Specifically, a stronger belief about a forthcoming cold stimulus resulted in heightened cold ratings (β = 0.15, 95% CI = [0.1; 0.19], p < .0001), but reduced warm ratings (β = −0.09, 95% CI = [−0.13; −0.05], p < .0001, Fig 4A and Supplementary Table 4A). Prediction uncertainty also exerted a notable influence on the perceived thermosensory quality of the TGI (Fig 4A and Supplementary Table 4B). Here, precise expectations of cold intensified cold ratings (β = 0.17, 95% CI = [0.11; 0.23], p < .0001) and reduced warm (β = −0.17, 95% CI = [−0.22; −0.11], p < .0001), but did not significantly influence burning ratings (β = −0.04, 95% CI = [−0.12; 0.03], p = 0.26) during the illusion. Conversely, precise expectations of warmth heightened both warm and burning sensations, accentuating both heat enhancement and illusory pain components of TGI perception.

**Fig 4.**
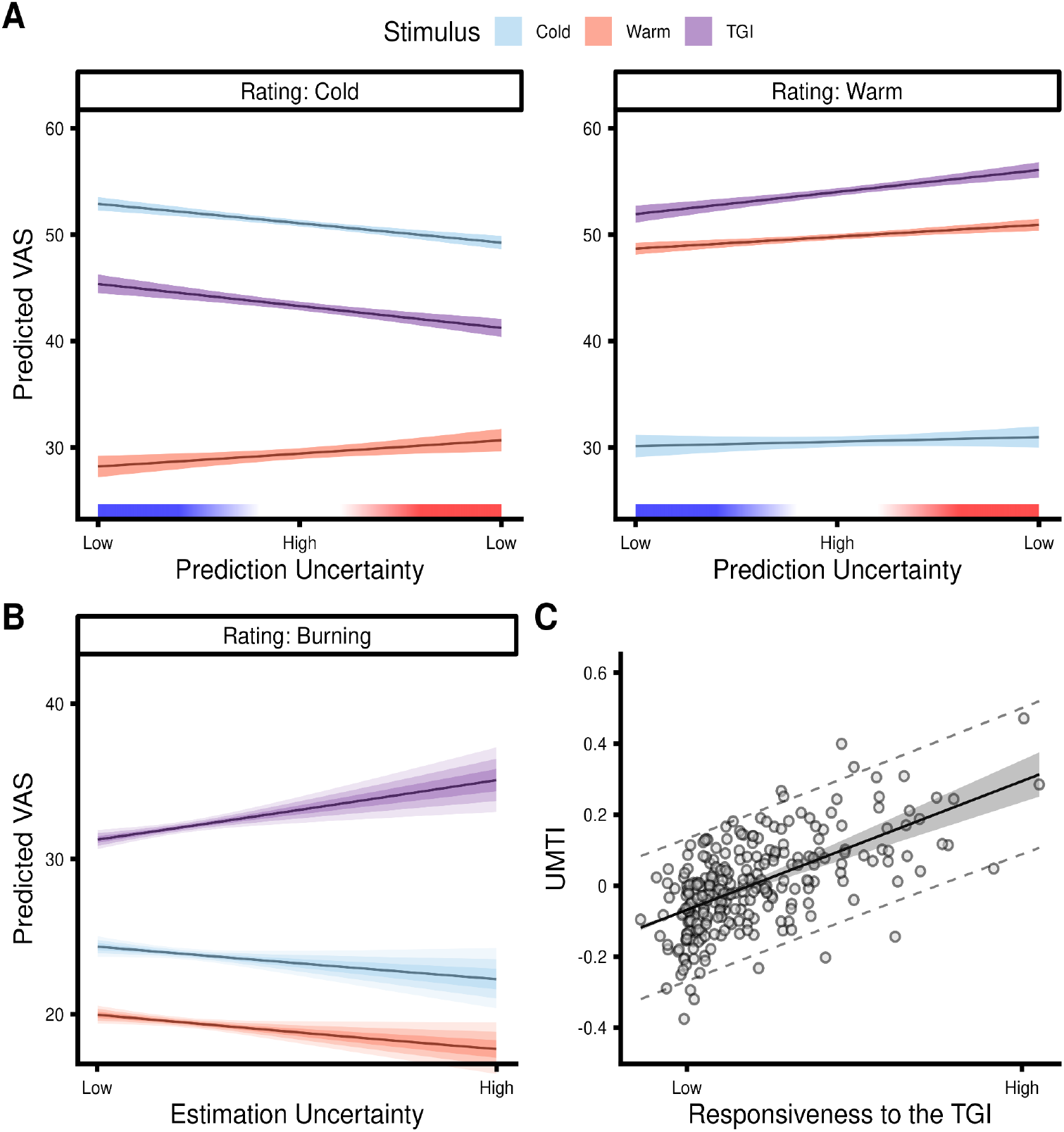
Effects of prediction and estimation uncertainty on veridical thermosensation and illusory pain. **A.** The impact of prediction uncertainty on thermosensory ratings for cold (blue), warm (red) and TGI stimuli (purple). The x-axis represents the precision of the lower-level belief about the forthcoming stimulus. Prediction uncertainty values range from high precision prediction that the stimulus would be warm to high precision predictions that the stimulus would be cold. Intermediate values indicate high prediction uncertainty about the thermal quality of the forthcoming stimulus. The y-axis indicates the predicted VAS ratings (i.e., marginal means) based on ZOIB modeling, separately for cold, warm and burning ratings with the shaded area depicting the 50, 80 and 95% confidence interval on the marginal means. **B.** The impact of estimation uncertainty on TGI perception. The x-axis depicts the varying degree of estimation uncertainty from low to high. The y-axis indicates the predicted burning ratings based on ZOIB modeling, separately for cold (blue), warm (red) and TGI stimuli (purple) with the shaded area depicting the 50, 80 and 95% confidence interval on the marginal means. **C.** Correlation between the TGI responsiveness (x-axis) and Uncertainty Modulation of TGI Index (UMTI). The positive correlation demonstrated that individuals with heightened responsiveness to TGI are also more susceptible to modulation of their illusion by uncertainty.

The higher-level estimation uncertainty played a more pronounced role in influencing burning sensations within TGI trials (Fig 4B and Supplementary Table 4C). Whereas precise cold expectations at the lower level were linked to reduced burning sensations, weak or unclear associations between cues and predicted stimuli increased burning ratings compared to both cold (β = −0.07, 95% CI = [−0.08; −0.05], p < .0001) and warm stimuli (β = −0.03, 95% CI = [−0.05; −0.01], p < .0001). This indicated that the illusory pain aspect of the TGI was most intense under conditions of ambiguous cue-stimulus mappings, or high estimation uncertainty. In essence, while lower-level prediction uncertainty predominantly determined whether the TGI was perceived as more cold or warm, the characteristic burning sensation of the TGI was markedly influenced by higher-level estimation uncertainty. These findings indicate how increased uncertainty regarding forthcoming stimulus temperatures can lead to a distorted perception of innocuous temperatures, manifesting as an aberrant sensation of pain.

#### Individual differences in thermosensory learning and TGI sensitivity

As participants exhibited large variability in their experience of the TGI (Fig 2A-C), we investigated how these variations manifested in thermosensory learning. To this end, we formulated two indices related to TGI responsiveness and susceptibility to uncertainty during thermosensory learning. To capture individual differences in response to TGI stimuli, we computed the discrepancy between burning ratings for TGI stimuli and the highest burning rating for either innocuous cold or warm stimuli. This approach yielded a continuous scale of TGI responsiveness, spanning from negative values indicative of TGI non-responders to positive values representing TGI responders. In addition, to quantify susceptibility to uncertainty, we defined a parameter (β), which corresponds to the degree to which individual burning ratings in TGI trials were influenced by estimation uncertainty (i.e., an Uncertainty Modulation of TGI Index or UMTI), thus reflecting individual differences in modulation of TGI intensity by higher order uncertainty (see Supplementary Note). Our analysis revealed a positive correlation between the TGI responsiveness index and UMTI (r(265) = 0.6 [0.51; 0.67] t = 12.1 p < .0001, Fig 4C), indicating that individuals with a heightened response to the TGI were also more susceptible to modulation of their illusion by uncertainty. This finding highlights the crucial link between individual variability in thermosensory learning and the subjective experience of TGI, and represents a novel, individually meaningful, metric of TGI responsiveness defined via computational modeling.

#### Computational fingerprints of thermosensory learning

We examined the neurobiological underpinnings of inter-individual variability in computational parameters, by relating ζ (capturing variability in the decision-making process, also known as decision temperature), ω (reflecting the speed of adaptation to changing conditions or learning) and UMTI (the uncertainty modulation of TGI index) parameters to brain microstructure. To this aim, we performed whole-brain voxel-based quantification analyses of Magnetization Transfer (MT), Longitudinal Relaxation Rate (R1) and effective transverse relaxation rate (R2*) weighted maps obtained from 0.8 mm Multi-Parameter Mapping (Table 1 and Fig 5). MT and R1 serve as indicators of cortical myeloarchitecture, aiding in the identification of myelination levels in gray matter. R2* is influenced by factors such as iron concentration [46,47]. A benefit of this approach is that the VBQ technique yields quantitative measures of local brain microstructure which are inherently meaningful and comparable across imaging sites or studies, unlike classical volumetric techniques which derive arbitrary signal units [47].

**Fig 5.**
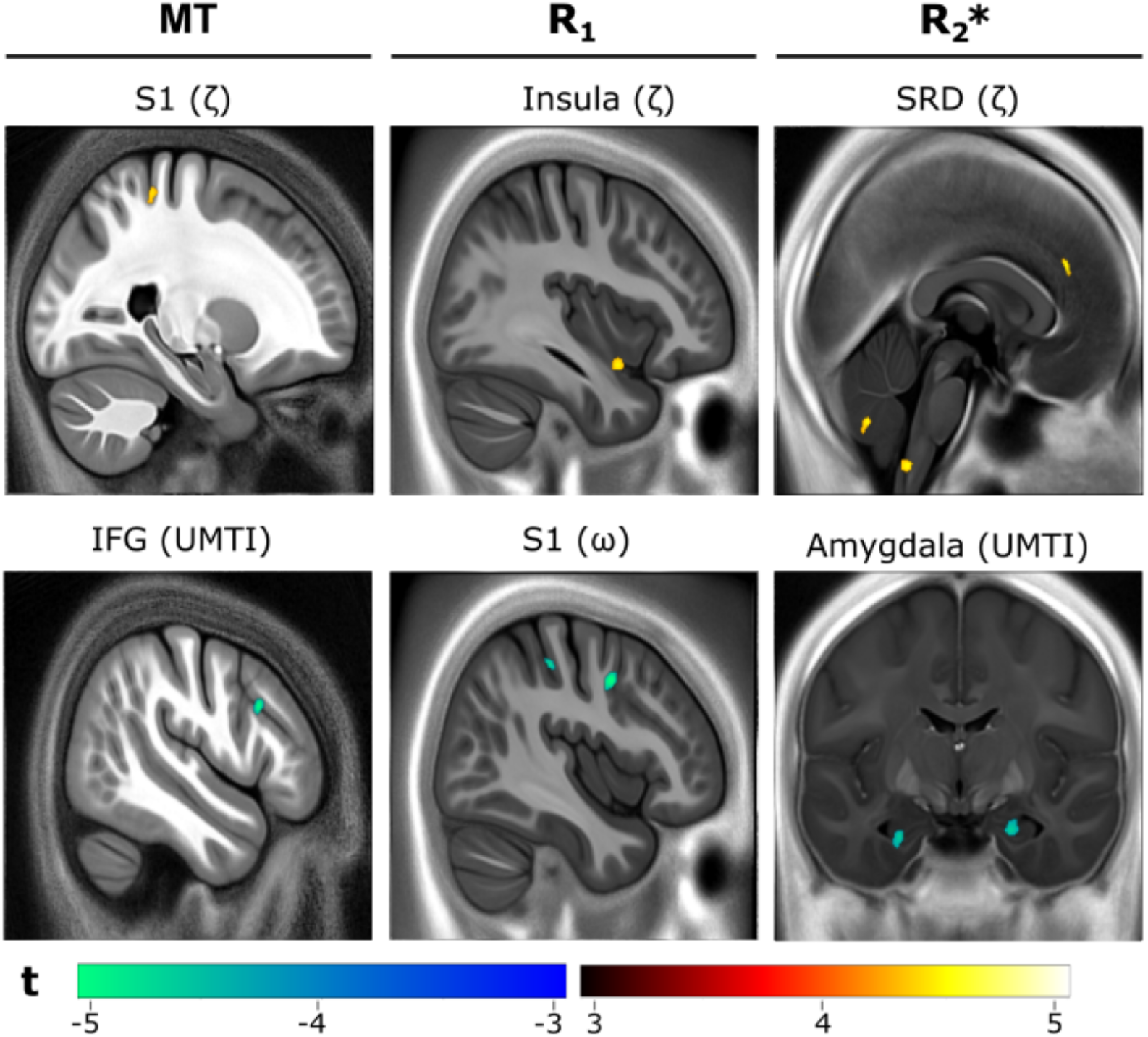
Microstructural brain correlates of computational parameters. Analysis of Multi-Parameter Maps indexing local gray matter myeloarchitecture (MT and R1) and iron content (R2*). Heat maps and color bars indicate voxelwise Z-statistics. For visualization purposes, thresholded Z-maps are plotted on the average normalized parametric map across the entire sample. Maps are family wise error (FWE) cluster-corrected for multiple comparisons, pFWE < 0.05. Columns represent MT, R1 and R2* maps respectively from left to right. Each cell denotes a specific map with names representing the anatomical identifier with the particular contrast delineated within parentheses.

**Table 1:** Microstructural brain correlates of computational parameters.

| **Region** | **k** | **p(FWE-corr)** | **Z value** | **x** | **y** | **z** | **MAP** | **Contrast** |
| --- | --- | --- | --- | --- | --- | --- | --- | --- |
| L Postcentral Gyrus | 210 | 0.005 | 3.84 | -27 | -39 | 63 | MT | $\zeta$ |
| R Inferior Frontal Gyrus | 606 | <.001 | -4.83 | 49 | 18 | 29 | MT | UMTI |
| R Postcentral Gyrus | 266 | 0.001 | -4.20 | 41 | -29 | 46 | R1 | $\omega$ |
| R Precentral Gyrus | 552 | <.001 | -5.28 | 43 | 3 | 40 | R1 | $\omega$ |
| R Posterior Insula | 226 | 0.003 | 3.77 | 38 | 5 | -16 | R1 | $\zeta$ |
| Subnucleus reticularis dorsalis | 202 | 0.02 | 3.70 | 1 | -45 | -58 | R2* | $\zeta$ |
| R Cerebellum Exterior | 396 | <.001 | 4.25 | 4 | -66 | -40 | R2* | $\zeta$ |
| L Superior Frontal Gyrus Medial Segment | 362 | <.001 | 4.23 | -5 | 38 | 30 | R2* | $\zeta$ |
| R Amygdala | 303 | 0.001 | -3.99 | 22 | -10 | -24 | R2* | UMTI |
| L Amygdala | 171 | 0.055 | -3.94 | -26 | -10 | -26 | R2* | UMTI |

We identified significant correlations between computational parameters and individual variation in the microstructure of brain regions previously related to pain and thermosensation (Table 1). Individual variability of decision-making processes, denoted by the parameter ζ, showed positive correlations with cortical myeloarchitecture in the left postcentral gyrus (MT map) and the right posterior insula (R1 map), as well as the iron concentration of the subnucleus reticularis dorsalis within the brainstem, among other regions such as the right cerebellum and the left superior frontal gyrus (R2* map). Additionally, the myeloarchitecture of the right posterior gyrus and the right precentral gyrus (R1 map) exhibited negative correlations with ω, highlighting the role of somatosensory and motor cortices in thermosensory associative learning. Finally, UMTI showed a negative correlation with the myeloarchitecture of the right inferior frontal gyrus (MT map) and with the iron concentration of the right basolateral amygdala (R2* map). This suggests that the influence of uncertainty on the perception of illusory pain is linked to the microstructural characteristics of regions involved in salience detection, as well as threat and fear learning [48]. This aligns with the view that inferior frontal gyrus and amygdala regions are instrumental in erroneously interpreting innocuous stimuli as painful, influenced by the brain’s estimation of uncertainty. These results shed light on the cortical myeloarchitecture fingerprints associated with ζ, ω, and UMTI, enhancing our understanding of the neurobiological underpinnings of decision-making, learning, and the modulation of uncertainty within the context of pain illusions. It also identifies structural features and regions that may potentially serve as biomarkers for these cognitive processes in individuals with chronic pain conditions.

## Discussion

We demonstrate that expectations shape both innocuous thermosensation and the Thermal Grill Illusion (TGI), providing a computational perspective on how the brain’s interpretation of innocuous thermal stimuli can paradoxically lead to pain. Through computational modeling, we have defined a mechanistic framework where hierarchical levels of predictions govern both the perceived quality of thermosensory inputs and the intensity of illusory pain. At the lower-level, the immediate predictions about an upcoming sensory stimulus modulate the perceived thermosensory quality. This modulation aligns closely with the predicted outcome, with the degree of influence scaled by the prediction’s uncertainty. In other words, more uncertain predictions result in less pronounced effects on temperature perception. At the higher level, the model encapsulates beliefs about cue-outcome associations, where greater uncertainty in these associations is found to amplify the experience of illusory pain, as evidenced by the increased burning sensations during the Thermal Grill Illusion (TGI).

We further define a computationally-based metric of TGI responsivity related to estimation uncertainty, referred to as Uncertainty Modulation of TGI Index (UMTI). We find that UMTI is positively correlated with TGI responsiveness, suggesting that illusory pain reflects increased susceptibility to thermosensory estimation uncertainty. This finding suggests that those who experience more pronounced illusory pain in the TGI tend to have a higher degree of uncertainty in discerning thermal sensations. Essentially, when their ability to anticipate and interpret thermal inputs is more uncertain, they are more prone to perceive these non-painful stimuli as painful.

We linked three distinct computational parameters of thermosensory learning and TGI perception to brain microstructure. Zeta (ζ), a parameter related to decision temperature, reflecting individual variability in the decision-making process was associated with gray matter myelination of primary somatosensory cortex and posterior insula, as well as iron content in the subnucleus reticularis dorsalis - a region in the medulla known for its involvement in the descending modulatory system [49,50]. Omega (ω), as a parameter reflecting the influence of uncertainty on learning, was also linked to the myelination of the primary somatosensory cortex. Finally, we identified novel brain markers inscribed in the iron concentration of basolateral amygdala and cortical myelination of the inferior frontal gyrus related to uncertainty modulation of TGI perception (UMTI). All these areas are notably involved in either pain processing [30,31] or threat detection [48], suggesting direct relationships between microstructure features in these regions and the computational mechanisms underlying illusory pain perception. Future research will focus on modeling functional connectivity within a brain network that includes these regions, using task-based fMRI imaging.

In summary, this study not only refines our understanding of human thermosensation from a Bayesian perspective but also identifies how uncertainty can lead to the misinterpretation of harmless stimuli as painful, transforming objectively innocuous stimuli into a subjective experience of pain. Our findings align with prior evidence supporting Bayesian models in pain perception and learning [23,24] and extend such framework to the domains of thermosensation and thermo-nociceptive illusions.

By showing that innocuous thermosensation and pain perception can be modulated by learned expectations, our study provides a computational framework that may be used to identify computational profiles in chronic pain conditions, where pain perception is decoupled from nociceptive inputs. For instance, in neuropathic pain conditions, often marked by nerve damage and resultant sensory disruption, individuals might experience heightened pain due to a misinterpretation of uncertain thermosensory inputs. Alternatively, chronic pain might be a result of the brain’s tendency to maintain its expectation of pain, which is not updated by sensory inputs due to the reduced sensory drive. These scenarios highlight how uncertainty and expectations can shape the experience of pain, as a result of altered bottom-up signaling. Overall, our study offers a computational framework to investigate how individual differences in learning could underlie experiences of chronic pain.

## Online methods

### Participants

A total of 273 participants completed a behavioral session of the probabilistic thermosensory learning task (PTL). We excluded six participants from the analyses due to missing responses in more than 10% of choices (i.e., predictions) or VAS ratings. The sample included in the behavioral analyses corresponded to 267 (182 female) participants, between the age of 18 and 52 years (mean = 24.5, sd = 4.4). 213 of these participants completed an MRI session, on a separate day, prior to the completion of the PTL task. Both MRI and behavioral sessions were completed within a three-week interval by the same individuals. This research, a subsection of a larger neuroimaging study with a total of 502 participants, involved various imaging, physiological, and cognitive assessments, focusing here on thermosensory learning and quantitative MRI data. Participants provided informed consent prior to the beginning of the study. The project received ethical approval from the Midtjylland Ethics Committee, and was conducted in accordance with the Declaration of Helsinki.

### Stimuli

Thermal stimuli were administered via a Thermal Cutaneous Stimulator (TCS) on the non-dominant forearm, allowing for quick and accurate responses with the dominant hand. The total stimulation surface covered 10^2^, comprising five distinct stimulation zones measuring 7 x 28 mm each. Cold and warm temperatures were individually calibrated, using a procedure that combined the method of limits and method of levels approaches. Innocuous warm stimuli involved three adjacent zones at an average temperature of 39.1 ± 2.8 °C while innocuous cold stimuli consisted of two adjacent zones at an average temperature of 20 ± 6.5 °C. The inactive zones remained at the baseline temperature of 32°C. TGI stimuli used the same temperatures as the innocuous conditions, with three warm and two cold stimuli presented in an alternating spatial pattern. Auditory tones, which served as cues for the forthcoming thermal stimulation, were either a lower tone of 400 Hz or a higher tone of 1600 Hz.

### Experimental procedure

In the PTL, participants completed 306 trials. Each trial began with a fixation point displayed for a random interval between one to two seconds. Following this, an auditory tone (either 400 or 1600 Hz) was presented, leading to a prompt for the participant to predict the upcoming thermosensory stimulus. This prediction was a binary choice - ‘cold’ or ‘warm’ - made using the left and right arrow keys, with a response time limit of three seconds. The stimulus, lasting 10 seconds, varied between cold (43% of trials), warm (43% of trials), or a combination of both in an alternating pattern to induce Thermal Grill Illusion (TGI), present in 14% of trials. In around 47% of trials, we collected Visual Analog Scale (VAS) ratings from participants to measure their perception of cold, warm, and burning sensations. Each sensation was rated on a separate scale ranging from 0 (no sensation) to 100 (maximum sensation), with these ratings required for approximately 60% of cold and warm trials and all TGI trials. Participants had up to five seconds to provide each VAS rating.

The likelihood of cue-stimulus associations was governed by two predetermined sequences, which were counterbalanced across participants. These sequences were designed to create blocks of stimuli where a specific cue had an 82% chance of indicating a particular outcome. However, these cues were subject to unpredictable reversals - at certain points, a cue that previously had an 82% likelihood of predicting one outcome would switch to having only an 18% likelihood of predicting that same outcome. Interspersed between these reversals were blocks of trials in which the association between a cue and a stimulus was at chance level.

### Statistical modeling

The modeling of error rates, response times and VAS ratings was conducted using generalized linear mixed effects models using the GAMLSS package [51] in R. Error rates were modeled using a binomial distribution with the logit link function, while response times were modeled using a gamma distribution with a logarithmic link function, VAS-ratings were modeled using the zero-one-inflated beta distribution with the logit link function for all parameters (see Supplementary Note). For all mixed effects models random intercepts were incorporated for each subject, incorporating random slopes where convergence permitted.

### Computational modeling

We compared three computational learning models: 2-level HGF, Rescorla Wagner and Sutton K1. To ensure the robustness of all models, we demonstrated acceptable parameter recovery, across a wide range of subject-specific learning parameters, as well as effective model recovery. Results from these analyses, further elaborated in Fig S4, ensured that the parameters derived from the models were interpretable and sensible, and facilitated the specification of reasonable parameter ranges for weakly informative priors for all models. For selecting the model that most accurately represented the data from our learning task, we employed a random effects model comparison using the VBA-toolbox [45]. This analysis indicated that the 2-level HGF was the most appropriate model to describe our data.

The 2-level HGF utilizes variational Bayesian approximation to derive update equations, enabling the estimation of how beliefs across different hierarchical levels evolve over trials [43,44]. The model is structured in two sub-parts known as response and perceptual models (Fig 3). The response model includes observed values, such as the inferred cue-outcome association (U), the participant’s prediction responses (Resp), and the decision temperature parameter zeta (ζ). The perceptual model is organized across two distinct hierarchical levels. At the first level (*x*_1_), the model captures participants’ immediate predictions about upcoming stimuli. These lower-level predictions follow a Bernouli distribution where the more extreme values around zero and one signify low prediction uncertainty, while intermediate values (0.5) indicate high uncertainty. The second level (*x*_2_) encapsulated predictions about the stability of cue-outcome associations. These higher-level beliefs evolve over time as a Gaussian random walk, with a step size determined by the parameter omega (ω).

The transition from the second-level to the first-level HGF is governed by a sigmoid transformation, converting the continuous Gaussian-distributed beliefs into discrete Bernoulli-distributed probabilities about immediate outcomes. The sigmoid transformation is formulated as:

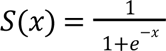

Simultaneously, the second-level HGF is updated based on precision-weighted prediction errors computed at the first-level. This update mechanism allows the model to account for confidence in the first-level predictions, adjusting the second-level beliefs accordingly. In cases of higher precision, when the uncertainty weight on prediction errors is low, the model is more likely to maintain its current belief structure. This belief updating is formulated as:

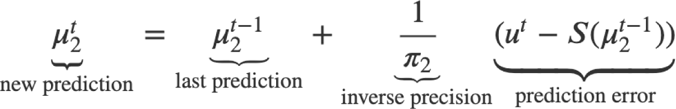

Where subscripts represent the level of the HGF, µ is the mean of the level, π is the precision of the level and *u* is the contingency input. In our computational analysis, this inverse precision is referred to as estimation uncertainty, while the first level uncertainty is labeled prediction uncertainty.

### Quantitative MRI

Quantitative MRI (qMRI) data were acquired using a 3T MRI scanner (Siemens Prisma), standard 32-channel RF head coil and a radiofrequency body coil. We obtained whole-brain images at isotropic 0.8mm resolution using a Multi-Parameter Mapping (MPM) quantitative imaging protocol [52]. The imaging sequences included three spoiled multi-echo 3D fast low angle shot (FLASH) acquisitions and three additional calibration sequences to correct for RF transmit field inhomogeneities (i.e., B1 mapping). Specifically, the FLASH sequences consisted of magnetization transfer (MT), proton density (PD) and T1 weighting acquisitions. The flip angle was 6° for MT and PD, while 21° for T1-weighted images. MT-weighting used a Gaussian RF pulse 2 kHz off resonance with 4ms duration and a nominal flip rate of 220°. The field of view was 256mm head-foot, 224mm anterior-posterior, and 179mm right-left. We acquired gradient echoes with alternating readout gradient polarity using equidistant echo times ranging from 2.34 to 13.8ms (MT) or 18.4ms (PD and T1), using a readout bandwidth of 490 Hz/pixel. For the MT-weighted acquisition, only 6 echoes were collected to achieve a repetition time (TR) of 25ms for all FLASH volumes. For accelerated data acquisition, we performed partially parallel imaging using the GRAPPA algorithm, with an acceleration factor of 2 in each phase encoded direction and 40 integrated reference lines. All acquisitions had a slab rotation of 30°. The B1 mapping acquisition comprised 11 measurements with the nominal flip rate ranging from 115° to 65° in 5° steps. The total scanning time for the qMRI acquisitions was approximately 26 minutes.

### Map creation

We obtained quantitative MT, PD, R1 and R2* maps using the hMRI toolbox v. 0.5.0 (January 2023) [53] and SMP12 (version 12.r7771, Wellcome Trust Centre for Neuroimaging, http://www.fil.ion.ucl.ac.uk/spm/). Except for enabling the correction for imperfect spoiling [54], the hMRI toolbox was configured using the standard settings. Prior to the estimation of these maps, all images were aligned to the MNI standard space. This processing produced four maps, each reflecting different attributes of brain tissue microstructure. The MT map is sensitive to myelination and myeloarchitecture, the PD map is indicative of tissue water content, the R1 map primarily reflects myelination, but also iron and water content, while the R2* map is sensitive to tissue iron concentration. We analyzed three of these maps (MT, R1 and R2*) independently.

We used the unified segmentation approach [55] to segment MT saturation maps into probability maps of gray matter (GM), white matter (WM) and cerebrospinal fluid (CSF). For this segmentation, we employed tissue probability maps based on multi-parameter data [56], without bias field correction as MT maps do not show significant bias field modulation. Subsequently, GM and WM probability maps were utilized for inter-subject registration using the nonlinear diffeomorphic algorithm DARTEL [57]. This step enabled the normalization of the derived quantitative maps to MNI space at an isotropic 1 mm resolution. This normalization used the DARTEL template created during registration and participant-specific deformation fields.

The nonlinear registration of the quantitative maps was based on the MT maps, chosen for their high contrast in subcortical structures and a WM-GM contrast in the cortex comparable to that of T1-weighted images [58]. Finally, tissue-weighted smoothing was applied using a 4 mm full width at half maximum (FWHM) kernel [59], to preserve quantitative values. The resulting smoothed, modulated and normalized GM images were used for statistical analyses. For visualization purposes, we generated average MT, R1 and R2* maps in standard space based on data from 442 individuals, who participated in the Visceral Mind Project. For analysis, we utilized a gray-matter mask that was generated by averaging the smoothed, modulated GM segments, and applying a threshold of p(gray matter) > 0.2.

### MPM Quality Control

We implemented a comprehensive set of quality-control (QC) protocols including manual and automated procedures. These included manual scoring of raw image quality at the time of acquisition, automated processing via the Mriqc pipeline [60], as well as the application of a specially developed semi-automatic hMRI-vQC pipeline designed for quantitative neuroimaging. We additionally calculated the index of motion degradation using QUQUI [61]. The automated procedures yielded a variety of quantitative QC metrics including coregistration parameters, standard deviation in the white matter of the R2* maps, SNR and CNR values; these were inspected via boxplots to identify extreme subjects. Following the application of MRIqc, a team of two authors (NN and CS) inspected all raw images and post-processed maps including those flagged by the automated procedure. Images were graded on a scale from 0-3 (unusable to no issues), and any disagreements between the raters were discussed and resolved alongside a third author (MA). In the Visceral Mind Project, 60 out of 502 individuals failed to pass these QC protocols, while in the subsample of the current study no participant was excluded from the voxel-based quantification analyses due to data quality.

### Voxel Based Quantification Analysis

We analyzed whole-brain associations between MT at each voxel and thermosensory learning using a multiple linear regression approach known as voxel-based quantification (VBQ). Our key analysis comprised positive and negative t-tests over the computational parameters. We included age, gender, and total intracranial volume as nuisance covariates in the regression model, following recommended procedures for computational neuroanatomy [62]. The analysis was adjusted for multiple comparisons using a cluster-based approach with non-stationarity correction, using a family wise error (FWE) cluster threshold of p < 0.025 (Bonferroni corrected for two one-tailed tests), based upon a p < 0.001 uncorrected inclusion threshold [62], within the gray matter mask. All statistical analyses were conducted in SPM12, while anatomical labels were determined using the JuBrain Anatomy Toolbox v. 3.0 [63]. The full suite of statistical maps generated from this study is available at a Neurovault repository.

## Data availability

Behavioral and modeling data are available at OSF. Brain imaging data are available at Neurovault.

## Code availability

All code is available at Github.

## Acknowledgements

This work was supported by the European Research Council under the grants ERC-2020-StG-948838 (FF, JFE, CEK, AGM, CSD) and ERC-2020-StG-948788 (MGA, ASC, MV). We also acknowledge funding from the Lundbeck Foundation under the grants R272-2017-4345 (MGA, LB, MB, NN). CS is supported by a Neuroscience Academy Denmark (NAD) fellowship. Additionally, CM is supported by Wellcome Mental Health Award 226776/Z/22/Z, Independent Research Fund Denmark (DFF) Research Project 3166-00158B, Carlsberg Foundation Young Researcher Fellowship CF21-0439, and Aarhus University Research Foundation Starting Grant AUFF-E-2019-7-10. We thank Tobias Hauser for comments on the study design and draft.

## Authors contributions

Author contributions listed alphabetically according to CRediT taxonomy: Conceptualization: MGA, FF. Data curation: MGA, DEC, JFE, FF, NN, CS. Formal analysis: MGA, DEC, JFE, FF, NN, CS. Funding acquisition: MGA, FF. Investigation: MGA, MB, DEC, JFE, FF, CEK, NN, MV Methodology: MGA, DEC, JFE, FF, CM. Project administration: MGA, FF, NN, MV. Resources: MGA, LB, MB, ASC, CSD, JFE, FF, CEK, AGM, CM, NN, MV. Software: MGA, DEC, JFE, FF, CM, NN. Supervision: MGA, FF. Visualization: MGA, JFE, FF. Writing – original draft: MGA, JFE, FF. Writing – review & editing: MGA, ASC, JFE, FF, CM, AGM, NN.

## References

1. Caterina MJ, Schumacher MA, Tominaga M, Rosen TA, Levine JD, Julius D. The capsaicin receptor: A heat-activated ion channel in the pain pathway. Nature. 1997;389: 816–824. doi:10.1038/39807

2. McKemy DD, Neuhausser WM, Julius D. Identification of a cold receptor reveals a general role for TRP channels in thermosensation. Nature. 2002;416: 52–58. doi:10.1038/nature719

3. Peier AM, Moqrich A, Hergarden AC, Reeve AJ, Andersson DA, Story GM, et al. A TRP channel that senses cold stimuli and menthol. Cell. 2002;108: 705–715. doi:10.1016/s0092-8674(02)00652-9

4. Atlas LY, Bolger N, Lindquist MA, Wager TD. Brain Mediators of Predictive Cue Effects on Perceived Pain. Journal of Neuroscience. 2010;30: 12964–12977. doi:10.1523/JNEUROSCI.0057-10.2010

5. Fields HL. How expectations influence pain. PAIN. 2018;159 Suppl 1: S3–S10. doi:10.1097/j.pain.0000000000001272

6. Jepma M, Koban L, Doorn J van, Jones M, Wager TD. Behavioural and neural evidence for self-reinforcing expectancy effects on pain. Nature Human Behaviour. 2018;2: 838–855. doi:10.1038/s41562-018-0455-8

7. Nickel MM, Tiemann L, Hohn VD, May ES, Gil Ávila C, Eippert F, et al. Temporal–spectral signaling of sensory information and expectations in the cerebral processing of pain. Proceedings of the National Academy of Sciences. 2022;119: e2116616119. doi:10.1073/pnas.2116616119

8. Craig AD, Bushnell MC. The thermal grill illusion: Unmasking the burn of cold pain. Science. 1994;265: 252–255. doi:10.1126/science.8023144

9. Craig AD, Reiman EM, Evans A, Bushnell MC. Functional imaging of an illusion of pain. Nature. 1996;384: 258–260. doi:10.1038/384258a0

10. Fardo F, Beck B, Allen M, Finnerup NB. Beyond Labeled Lines: A Population Coding Account of the Thermal Grill Illusion. Neuroscience and Biobehavioral Reviews. 2020;108: 472–479. doi:10.1016/j.neubiorev.2019.11.017

11. Mitchell AG, Ehmsen JF, Christensen DE, Stuckert A, Haggard P, Fardo F. Disentangling the spinal mechanisms of illusory heat and burning sensations in the Thermal Grill Illusion. bioRxiv; 2023. doi:10.1101/2023.08.24.554485

12. Fardo F, Finnerup NB, Haggard P. Organization of the Thermal Grill Illusion by Spinal Segments. Annals of Neurology. 2018;84: 463–472. doi:10.1002/ana.25307

13. Büchel C, Geuter S, Sprenger C, Eippert F. Placebo analgesia: A predictive coding perspective. Neuron. 2014;81: 1223–1239. doi:10.1016/j.neuron.2014.02.042

14. Fardo F, Auksztulewicz R, Allen M, Dietz MJ, Roepstorff A, Friston KJ. Expectation violation and attention to pain jointly modulate neural gain in somatosensory cortex. NeuroImage. 2017;153: 109–121. doi:10.1016/j.neuroimage.2017.03.041

15. Geuter S, Boll S, Eippert F, Büchel C. Functional dissociation of stimulus intensity encoding and predictive coding of pain in the insula. Johansen-Berg H, editor. eLife. 2017;6: e24770. doi:10.7554/eLife.24770

16. Tabor A, Thacker MA, Moseley GL, Körding KP. Pain: A Statistical Account. PLOS Computational Biology. 2017;13: e1005142. doi:10.1371/journal.pcbi.1005142

17. Hoskin R, Berzuini C, Acosta-Kane D, El-Deredy W, Guo H, Talmi D. Sensitivity to pain expectations: A Bayesian model of individual differences. Cognition. 2019;182: 127–139. doi:10.1016/j.cognition.2018.08.022

18. Seymour B, Mancini F. Hierarchical models of pain: Inference, information-seeking, and adaptive control. NeuroImage. 2020;222: 117212. doi:10.1016/j.neuroimage.2020.117212

19. Song Y, Yao M, Kemprecos H, Byrne A, Xiao Z, Zhang Q, et al. Predictive coding models for pain perception. Journal of Computational Neuroscience. 2021;49: 107–127. doi:10.1007/s10827-021-00780-x

20. Eckert A-L, Pabst K, Endres DM. A Bayesian model for chronic pain. Frontiers in Pain Research. 2022;3. Available: https://www.frontiersin.org/articles/10.3389/fpain.2022.966034

21. Kiverstein J, Kirchhoff MD, Thacker M. An Embodied Predictive Processing Theory of Pain Experience. Review of Philosophy and Psychology. 2022;13: 973–998. doi:10.1007/s13164-022-00616-2

22. Chen ZS, Wang J. Pain, from perception to action: A computational perspective. iScience. 2023;26: 105707. doi:10.1016/j.isci.2022.105707

23. Mancini F, Zhang S, Seymour B. Computational and neural mechanisms of statistical pain learning. Nature Communications. 2022;13: 6613. doi:10.1038/s41467-022-34283-9

24. Mulders D, Seymour B, Mouraux A, Mancini F. Confidence of probabilistic predictions modulates the cortical response to pain. Proceedings of the National Academy of Sciences. 2023;120: e2212252120. doi:10.1073/pnas.2212252120

25. Roy M, Shohamy D, Daw N, Jepma M, Wimmer GE, Wager TD. Representation of aversive prediction errors in the human periaqueductal gray. Nature Neuroscience. 2014;17: 1607–1612. doi:10.1038/nn.3832

26. Fazeli S, Büchel C. Pain-Related Expectation and Prediction Error Signals in the Anterior Insula Are Not Related to Aversiveness. The Journal of Neuroscience: The Official Journal of the Society for Neuroscience. 2018;38: 6461–6474. doi:10.1523/JNEUROSCI.0671-18.2018

27. Levine JD, Gordon NC, Fields HL. The mechanism of placebo analgesia. Lancet. 1978;2: 654–657. doi:10.1016/s0140-6736(78)92762-9

28. Wager TD, Rilling JK, Smith EE, Sokolik A, Casey KL, Davidson RJ, et al. Placebo-induced changes in FMRI in the anticipation and experience of pain. Science. 2004;303: 1162–1167. doi:10.1126/science.1093065

29. Price DD, Finniss DG, Benedetti F. A comprehensive review of the placebo effect: Recent advances and current thought. Annual Review of Psychology. 2008;59: 565–590. doi:10.1146/annurev.psych.59.113006.095941

30. Eippert F, Bingel U, Schoell ED, Yacubian J, Klinger R, Lorenz J, et al. Activation of the opioidergic descending pain control system underlies placebo analgesia. Neuron. 2009;63: 533–543. doi:10.1016/j.neuron.2009.07.014

31. Eippert F, Finsterbusch J, Bingel U, Büchel C. Direct evidence for spinal cord involvement in placebo analgesia. Science. 2009;326: 404. doi:10.1126/science.1180142

32. Benedetti F. Placebo effects: From the neurobiological paradigm to translational implications. Neuron. 2014;84: 623–637. doi:10.1016/j.neuron.2014.10.023

33. Wager TD, Atlas LY. The neuroscience of placebo effects: Connecting context, learning and health. Nature Reviews Neuroscience. 2015;16: 403–418. doi:10.1038/nrn3976

34. Anchisi D, Zanon M. A Bayesian perspective on sensory and cognitive integration in pain perception and placebo analgesia. PloS One. 2015;10: e0117270. doi:10.1371/journal.pone.0117270

35. Tabor A, Burr C. Bayesian Learning Models of Pain: A Call to Action. Current Opinion in Behavioral Sciences. 2019;26: 54–61. doi:10.1016/j.cobeha.2018.10.006

36. Ongaro G, Kaptchuk TJ. Symptom perception, placebo effects, and the Bayesian brain. PAIN. 2019;160: 1–4. doi:10.1097/j.pain.0000000000001367

37. Craig AD. Can the basis for central neuropathic pain be identified by using a thermal grill? PAIN. 2008;135: 215–216. doi:10.1016/j.pain.2008.01.022

38. Adam F, Jouët P, Sabaté J-M, Perrot S, Franchisseur C, Attal N, et al. Thermal grill illusion of pain in patients with chronic pain: A clinical marker of central sensitization? PAIN. 2023;164: 638–644. doi:10.1097/j.pain.0000000000002749

39. Ouden HEM den, Daunizeau J, Roiser J, Friston KJ, Stephan KE. Striatal prediction error modulates cortical coupling. Journal of Neuroscience. 2010;30: 3210–3219. doi:10.1523/JNEUROSCI.4458-09.2010

40. Iglesias S, Mathys C, Brodersen KH, Kasper L, Piccirelli M, Ouden HEM den, et al. Hierarchical prediction errors in midbrain and basal forebrain during sensory learning. Neuron. 2013;80: 519–530. doi:10.1016/j.neuron.2013.09.009

41. Berker AO de, Rutledge RB, Mathys C, Marshall L, Cross GF, Dolan RJ, et al. Computations of uncertainty mediate acute stress responses in humans. Nature Communications. 2016;7: 10996. doi:10.1038/ncomms10996

42. Lawson RP, Mathys C, Rees G. Adults with autism overestimate the volatility of the sensory environment. Nature Neuroscience. 2017;20: 1293–1299. doi:10.1038/nn.4615

43. Mathys C, Daunizeau J, Friston K, Stephan K. A Bayesian Foundation for Individual Learning Under Uncertainty. Frontiers in Human Neuroscience. 2011;5. Available: https://www.frontiersin.org/articles/10.3389/fnhum.2011.00039

44. Mathys CD, Lomakina EI, Daunizeau J, Iglesias S, Brodersen KH, Friston KJ, et al. Uncertainty in perception and the Hierarchical Gaussian Filter. Frontiers in Human Neuroscience. 2014;8. Available: https://www.frontiersin.org/articles/10.3389/fnhum.2014.00825

45. Daunizeau J, Adam V, Rigoux L. VBA: A Probabilistic Treatment of Nonlinear Models for Neurobiological and Behavioural Data. PLOS Computational Biology. 2014;10: e1003441. doi:10.1371/journal.pcbi.1003441

46. Samson RS, Ciccarelli O, Kachramanoglou C, Brightman L, Lutti A, Thomas DL, et al. Tissue- and column-specific measurements from multi-parameter mapping of the human cervical spinal cord at 3 T. NMR in Biomedicine. 2013;26: 1823–1830. doi:10.1002/nbm.3022

47. Weiskopf N, Mohammadi S, Lutti A, Callaghan MF. Advances in MRI-based computational neuroanatomy: From morphometry to in-vivo histology. Current Opinion in Neurology. 2015;28: 313. doi:10.1097/WCO.0000000000000222

48. Fox AS, Oler JA, Tromp DPM, Fudge JL, Kalin NH. Extending the amygdala in theories of threat processing. Trends in neurosciences. 2015;38: 319–329. doi:10.1016/j.tins.2015.03.002

49. Mills EP, Di Pietro F, Alshelh Z, Peck CC, Murray GM, Vickers ER, et al. Brainstem Pain-Control Circuitry Connectivity in Chronic Neuropathic Pain. The Journal of Neuroscience: The Official Journal of the Society for Neuroscience. 2018;38: 465–473. doi:10.1523/JNEUROSCI.1647-17.2017

50. Youssef AM, Macefield VG, Henderson LA. Pain inhibits pain; human brainstem mechanisms. NeuroImage. 2016;124: 54–62. doi:10.1016/j.neuroimage.2015.08.060

51. Rigby RA, Stasinopoulos DM. Generalized Additive Models for Location, Scale and Shape. Journal of the Royal Statistical Society Series C (Applied Statistics). 2005;54: 507–554. Available: https://www.jstor.org/stable/3592732

52. Weiskopf N, Suckling J, Williams G, Correia MM, Inkster B, Tait R, et al. Quantitative multi-parameter mapping of R1, PD*, MT, and R2* at 3T: A multi-center validation. Frontiers in Neuroscience. 2013;7: 95. doi:10.3389/fnins.2013.00095

53. Tabelow K, Balteau E, Ashburner J, Callaghan MF, Draganski B, Helms G, et al. hMRI—A toolbox for quantitative MRI in neuroscience and clinical research. NeuroImage. 2019;194: 191–210. doi:10.1016/j.neuroimage.2019.01.029

54. Corbin N, Callaghan MF. Imperfect spoiling in variable flip angle T1 mapping at 7T: Quantifying and minimizing impact. Magnetic Resonance in Medicine. 2021;86: 693–708. doi:10.1002/mrm.28720

55. Ashburner J, Friston KJ. Unified segmentation. NeuroImage. 2005;26: 839–851. doi:10.1016/j.neuroimage.2005.02.018

56. Lorio S, Fresard S, Adaszewski S, Kherif F, Chowdhury R, Frackowiak RS, et al. New tissue priors for improved automated classification of subcortical brain structures on MRI. NeuroImage. 2016;130: 157–166. doi:10.1016/j.neuroimage.2016.01.062

57. Ashburner J. A fast diffeomorphic image registration algorithm. NeuroImage. 2007;38: 95–113. doi:10.1016/j.neuroimage.2007.07.007

58. Helms G, Draganski B, Frackowiak R, Ashburner J, Weiskopf N. Improved segmentation of deep brain grey matter structures using magnetization transfer (MT) parameter maps. Neuroimage. 2009;47: 194–198. doi:10.1016/j.neuroimage.2009.03.053

59. Draganski B, Ashburner J, Hutton C, Kherif F, Frackowiak RSJ, Helms G, et al. Regional specificity of MRI contrast parameter changes in normal ageing revealed by voxel-based quantification (VBQ). NeuroImage. 2011;55: 1423–1434. doi:10.1016/j.neuroimage.2011.01.052

60. Esteban O, Birman D, Schaer M, Koyejo OO, Poldrack RA, Gorgolewski KJ. MRIQC: Advancing the automatic prediction of image quality in MRI from unseen sites. PLOS ONE. 2017;12: e0184661. doi:10.1371/journal.pone.0184661

61. Lutti A, Corbin N, Ashburner J, Ziegler G, Draganski B, Phillips C, et al. Restoring statistical validity in group analyses of motion-corrupted MRI data. Human Brain Mapping. 2022;43: 1973–1983. doi:10.1002/hbm.25767

62. Ridgway GR, Henley SMD, Rohrer JD, Scahill RI, Warren JD, Fox NC. Ten simple rules for reporting voxel-based morphometry studies. NeuroImage. 2008;40: 1429–1435. doi:10.1016/j.neuroimage.2008.01.003

63. Eickhoff SB, Stephan KE, Mohlberg H, Grefkes C, Fink GR, Amunts K, et al. A new SPM toolbox for combining probabilistic cytoarchitectonic maps and functional imaging data. NeuroImage. 2005;25: 1325–1335. doi:10.1016/j.neuroimage.2004.12.034

