## Supplementary Material for "Uncertainty in Thermosensory Expectations Enhances an Illusion of Pain"

***Supplementary Results***

We investigated how response times were affected in trials subsequent to a thermal grill stimulus. Our findings revealed a significant influence of the interaction between cue-stimulus association and participants' perception of TGI quality ($\beta = 0.14$, 95% CI = [0.06; 0.21], $p < .0001$). Specifically, when there was a congruence between the predicted temperature (contingency) and the actual perceived TGI quality (e.g., anticipating cold and perceiving the TGI as predominantly cold), participants' response times on the trial following a TGI stimulus remained unchanged, indicating no post-TGI slowing ($\beta = -0.02$, 95% CI = [-0.07; 0.03], $p = 0.43$). Conversely, when there was a mismatch between the predicted temperature and perceived TGI quality (for instance, expecting warm but perceiving TGI as predominantly cold), participants exhibited slower response times in the subsequent trial ($\beta = 0.12$, 95% CI = [0.06; 0.17], $p < .0001$). Further details can be found in the supplementary table 2f).

***Supplementary Note***

**Formulation of reported models**

We modeled three types of responses: (1) binary choices, which determined if a participant predicted a cold or a warm stimulus, (2) response times associated with these binary choices and (3) VAS ratings, which reflected how a received stimulus was perceived by a participant. Here, we detail the probability distribution of each response type, as well as the parameters upon which our regression analysis is based. To model the binary choices, we used the binomial distribution which is given by:

$$f(y|\mu) = \frac{\Gamma(n+1)}{\Gamma(y+1)\Gamma(n-y+1)}\mu^{y}(1 - \mu)^{(n-y)}$$

Where Γ is the gamma function, y is the random variable of n successes (restricted to integer values) and $\mu \in [0, 1]$ is the probability of a given success. Here we parameterize μ using the logit link function (the inverse sigmoid transformation) $logit(\mu) = log\left(\frac{\mu}{1-\mu}\right)$.

To model response times, we used the gamma distribution given by:

$$f(y|\mu, \sigma) = \frac{y^{(1/\sigma^2-1)}e^{[-y/(\sigma^2\mu)]}}{(\sigma^2\mu)^{(1/\sigma^2)}\Gamma(1/\sigma^2)}$$

Where Γ is the gamma function, y is the random variable of response times (restricted to positive values), $\mu \in ]0, inf[$ is the mean of the distribution and $\sigma \in ]0, inf[$ is the square root of the usual dispersion parameter for a GLM gamma model. $\sigma * \mu$ is the standard deviation of the defined distribution. Here we parameterize μ using the logarithmic link function.

To model Visual analog scale (VAS) ratings, we used the zero one inflated beta (ZOIB) distribution, which is a mixture of two Bernoulli distributions and one beta distribution, formally given by:

$$beinf\left(y|p_0, p_1, \mu, \phi\right) = \{p_0\, y = 0\; f(y|\mu, \phi)\; y \in [0, 1]\; p_1\, y = 1$$

where the probability density function of the beta distribution $f(y; \mu, \phi)$ is given by

$$f(y; \mu, \phi) = \frac{\Gamma(\alpha+\beta)}{\Gamma(\alpha)\Gamma(\beta)}y^{\alpha-1}(1 - y)^{\beta-1}, y \in [0, 1]$$

In the GAMLSS packages, the parameters are parameterized as follows:

$$\mu = \frac{\alpha}{\alpha+\beta}$$

$$\sigma = \frac{1}{\alpha+\beta+1}$$

$$\tau = \frac{p_0}{p_2}$$

$$\nu = \frac{p_1}{p_2}$$

where $p_2 = 1 - p_0 - p_1$. All these given parameters μ, σ, τ and ν are restricted between 0 and 1, and are modeled using the logit link function.

**Formulation of the Uncertainty Modulation of TGI Index**

To provide a thorough understanding of the subject specific Uncertainty Modulation index parameter (UMTI), here we present the detailed mathematical formulation of the model. This formulation is written using the lmer syntax, as detailed below.

$$Burning_i \sim Est_i * Stim_i + Trial_i + (Est * Stim|ID),$$

$$family = beinf\left(\mu_i, \tau_i, \nu_i, \kappa\right)$$

Considering the number of parameters that have been parameterized, our primary focus in this section is on the mean. However, it is important to note that this approach is equally applicable to the parameters representing the proportion of ones and zeros (i.e., ν & τ). The mathematical description, specifically tailored to address only the mean, is as follows:

$$Burning_i \sim beinf\left(\mu_i, \tau, \nu, \kappa\right)$$

$$u_{i,j} = \beta_{0_J} + \beta_4 * Est_{i_j} + \beta_5 * Stim_{i_j} + \beta_6 * Trial_i + \beta_{1_J} * \left(Est_{i_j} * Stim(cold)_{i_j}\right) +$$

$$\beta_{2_J} * \left(Est_{i_j} * Stim(warm)_{i_j}\right) + \beta_{3_J} * \left(Est_{i_j} * Stim(TGI)_{i_j}\right)$$

Now, we present the structure of the random effects, illustrated through the variance-covariance matrix. Here, we exclude the upper triangle of the matrix to avoid redundancy.

$$\begin{pmatrix}\beta_{0_J}\\ \beta_{1_J}\\ \beta_{2_J}\\ \beta_{3_J}\end{pmatrix} \sim \mathcal{N}\left(\begin{pmatrix}\mu_{\beta_0}\\ \mu_{\beta_1}\\ \mu_{\beta_2}\\ \mu_{\beta_3}\end{pmatrix}, \begin{bmatrix}\sigma_{\beta_0} & . & . & .\\ \sigma_{\beta_0}\sigma_{\beta_1}\rho_1 & \sigma_{\beta_1} & . & .\\ \sigma_{\beta_0}\sigma_{\beta_2}\rho_2 & \sigma_{\beta_1}\sigma_{\beta_2}\rho_4 & \sigma_{\beta_2} & .\\ \sigma_{\beta_0}\sigma_{\beta_3}\rho_3 & \sigma_{\beta_1}\sigma_{\beta_3}\rho_5 & \sigma_{\beta_2}\sigma_{\beta_3}\rho_6 & \sigma_{\beta_3}\end{bmatrix}\right)$$

In this analysis, the parameter estimate of interest (i.e., UMTI) is $\beta_{3_j}$, which is the beta estimate for the j-th participant ID. This estimate specifically denotes the interaction term, which quantifies the degree to which estimation uncertainty influences the participant's response to the TGI, compared to their response to cold and warm stimuli. Positive values of $\beta_{3_j}$ suggest that a participant exhibits an increased tendency to rate the sensation as more 'burning' under TGI stimulus conditions, relative to either cold or warm stimuli, as estimation uncertainty increases. It is important to note that this effect is distinct from the direct stimulus effect of the TGI; it represents the differential impact of estimation uncertainty on burning ratings across stimulus types.

***Supplementary Tables***

**Table 1a,** Main effect of expectations on accuracy in predicting the next stimulus.

| predAcc ~ Expectation + Trial Number + re(random = ~ Trial number \| id), Binomial(link = logit) | | | | | |
|---|---|---|---|---|---|
| **parameter** | **contrast** | **β** | **SE** | **t** | **p** |
| μ | Intercept | 1.2 | 0.018 | 67 | 0 |
| μ | Expectation(Neutral) | -1.3 | 0.021 | -64 | 0 |
| μ | Expectation(Unpredicted) | -2.3 | 0.025 | -91 | 0 |
| μ | Trial Number | -0.00089 | 9.8e-05 | -9.1 | 7.5e-20 |

**Table 1b,** Main effect of expectations on response times in the next stimulus.

| predRT2 ~ Expectation + Trial number + re(random = ~1 \| id), Gamma(link = log) | | | | | |
|---|---|---|---|---|---|
| **parameter** | **contrast** | **β** | **SE** | **t** | **p** |
| μ | Intercept | -0.12 | 0.0042 | -28 | 2.5e-170 |
| μ | Expectation(Neutral) | 0.15 | 0.0059 | 25 | 4e-136 |
| μ | Expectation(TGI) | 0.34 | 0.0055 | 61 | 0 |
| μ | Expectation(Unpredicted) | 0.31 | 0.0056 | 56 | 0 |
| μ | Trial Number | -0.00034 | 2.3e-05 | -15 | 8e-52 |
| σ | Intercept | -0.55 | 0.005 | -1.1e+02 | 0 |
| σ | Expectation(Neutral) | -0.042 | 0.0068 | -6.2 | 7.7e-10 |
| σ | Expectation(TGI) | -0.21 | 0.0072 | -29 | 7.3e-184 |
| σ | Expectation(Unpredicted) | -0.18 | 0.0072 | -25 | 3.1e-141 |
| σ | Trial Number | 0.00036 | 2.7e-05 | 13 | 4.2e-40 |

**Table 2a,** Main effect of TGI vs. cold and warm on burning ratings.

| BurnBeta ~ Stimulus + Trial Number + re(random = ~ Stimulus \| id), ZOIB(link = logit) | | | | | |
|---|---|---|---|---|---|
| **parameter** | **contrast** | **β** | **SE** | **t** | **p** |
| μ | Intercept | -1 | 0.013 | -79 | 0 |
| μ | Stimulus(Cold) | -0.45 | 0.012 | -37 | 1.8e-290 |
| μ | Stimulus(Warm) | -0.65 | 0.012 | -53 | 0 |
| μ | Trial Number | 0.00068 | 5.7e-05 | 12 | 2.4e-32 |
| σ | Intercept | -0.6 | 0.014 | -43 | 0 |
| σ | Stimulus(Cold) | 0.014 | 0.013 | 1 | 0.3 |
| σ | Stimulus(Warm) | -0.057 | 0.013 | -4.3 | 2.1e-05 |
| σ | Trial Number | -0.00017 | 6.1e-05 | -2.7 | 0.0069 |
| ν | Intercept | -1.2 | 0.082 | -14 | 1.2e-45 |
| ν | Stimulus(Cold) | 2.2 | 0.077 | 29 | 5.7e-184 |
| ν | Stimulus(Warm) | 3.5 | 0.076 | 47 | 0 |
| ν | Trial Number | -0.00046 | 0.00034 | -1.3 | 0.18 |
| τ | Intercept | -6.1 | 0.27 | -23 | 3.9e-114 |
| τ | Stimulus(Cold) | -1.6 | 0.25 | -6.4 | 1.5e-10 |
| τ | Stimulus(Warm) | -1.6 | 0.25 | -6.5 | 1.1e-10 |
| τ | Trial Number | 0.0062 | 0.0012 | 5.2 | 2.4e-07 |

**Table 2b,** Main effect of stimulus on cold ratings.

| **ColdBeta ~ Stimulus + Trial Number + re(random = ~ Stimulus \| id), ZOIB(link = logit)** | | | | | |
|---|---|---|---|---|---|
| **parameter** | **contrast** | **β** | **SE** | **t** | **p** |
| μ | Intercept | -0.17 | 0.009 | -19 | 9.6e-83 |
| μ | Stimulus(TGI) | -0.39 | 0.0089 | -44 | 0 |
| μ | Stimulus(Warm) | -1 | 0.011 | -88 | 0 |
| μ | Trial Number | -0.00047 | 4.5e-05 | -10 | 9.4e-26 |
| σ | Intercept | -0.74 | 0.012 | -62 | 0 |
| σ | Stimulus(TGI) | 0.13 | 0.012 | 11 | 4.5e-26 |
| σ | Stimulus(Warm) | 0.44 | 0.013 | 35 | 7e-258 |
| σ | Trial Number | -0.00025 | 5.7e-05 | -4.4 | 1e-05 |
| ν | Intercept | -5.4 | 0.088 | -61 | 0 |
| ν | Stimulus(TGI) | 2.2 | 0.078 | 28 | 1.1e-176 |
| ν | Stimulus(Warm) | 6.6 | 0.088 | 75 | 0 |
| ν | Trial number | 0.0028 | 0.00035 | 7.9 | 2.3e-15 |
| τ | Intercept | -5.5 | 0.23 | -24 | 3e-123 |
| τ | Stimulus(TGI) | -0.76 | 0.25 | -3 | 0.0025 |
| τ | Stimulus(Warm) | -0.85 | 0.27 | -3.2 | 0.0014 |
| τ | Trial Number | -0.0013 | 0.0012 | -1.1 | 0.28 |

**Table 2c,** Main effect of stimulus on warm ratings.

| **Warm(Beta) ~ Stimulus + Trial Number + re(random = ~ Stimulus \| id), ZOIB(link = logit)** | | | | | |
|---|---|---|---|---|---|
| **parameter** | **contrast** | **β** | **SE** | **t** | **p** |
| μ | Intercept | -0.34 | 0.0084 | -40 | 0 |
| μ | Stimulus(Cold) | -0.82 | 0.011 | -72 | 0 |
| μ | Stimulus (TGI) | 0.18 | 0.0083 | 22 | 5.2e-108 |
| μ | Trial Number | 0.00023 | 4.2e-05 | 5.4 | 5.7e-08 |
| σ | Intercept | -0.8 | 0.011 | -70 | 0 |
| σ | Stimulus(Cold) | 0.53 | 0.013 | 42 | 0 |
| σ | Stimulus (TGI) | 0.2 | 0.011 | 17 | 2.4e-67 |
| σ | Trial Number | -0.00043 | 5.5e-05 | -7.7 | 1e-14 |
| ν | Intercept | -4.8 | 0.083 | -58 | 0 |
| ν | Stimulus(Cold) | 5.9 | 0.082 | 72 | 0 |
| ν | Stimulus (TGI) | 1.3 | 0.081 | 16 | 1.2e-57 |
| ν | Trial Number | -0.00056 | 0.00035 | -1.6 | 0.11 |
| τ | Intercept | -6.6 | 0.26 | -26 | 2.5e-145 |
| τ | Stimulus(Cold) | -1.7 | 0.49 | -3.5 | 0.00046 |
| τ | Stimulus (TGI) | 1.3 | 0.22 | 6 | 1.9e-09 |
| τ | Trial Number | 0.00049 | 0.0012 | 0.42 | 0.68 |

**Table 2d,** Expectation effects of thermosensory ratings.

| **value ~ RateCon * Stimulus * Prediction + Trial Number + re(random = ~ Stimulus \| id), ZOIB(link = logit)** | | | | | |
|---|---|---|---|---|---|
| **parameter** | **contrast** | **β** | **SE** | **t** | **p** |
| μ | Intercept | -1.2 | 0.017 | -69 | 0 |
| μ | RatingScale(Factual) | 0.89 | 0.018 | 50 | 0 |
| μ | Stimulus(Cold) | 0.1 | 0.022 | 4.6 | 5e-06 |
| μ | Prediction(Cold) | 0.038 | 0.023 | 1.7 | 0.094 |
| μ | Trial Number | -6.5e-05 | 3.9e-05 | -1.7 | 0.097 |
| μ | RatingScale (Factual): Stimulus(Cold) | -0.11 | 0.025 | -4.5 | 5.6e-06 |
| μ | RatingScale (Factual): Prediction(Cold) | -0.086 | 0.025 | -3.5 | 0.00053 |
| μ | Stimulus (Cold): Prediction(Cold) | -0.099 | 0.032 | -3.1 | 0.0016 |
| μ | RatingScale(Factual): Stimulus (Cold): Prediction(Cold) | 0.24 | 0.035 | 6.8 | 9.1e-12 |
| σ | Intercept | -0.044 | 0.016 | -2.7 | 0.0075 |
| σ | RatingScale(Factual) | -0.69 | 0.018 | -39 | 2e-323 |
| σ | Stimulus(Cold) | -0.11 | 0.02 | -5.6 | 2.5e-08 |
| σ | Prediction(Cold) | -0.018 | 0.02 | -0.88 | 0.38 |
| σ | Trial Number | -0.00037 | 4.9e-05 | -7.6 | 4e-14 |
| σ | RatingScale (Factual): Stimulus(Cold) | 0.25 | 0.026 | 9.7 | 3e-22 |
| σ | RatingScale (Factual): Prediction(Cold) | 0.0071 | 0.025 | 0.28 | 0.78 |
| σ | Stimulus (Cold): Prediction(Cold) | 0.026 | 0.029 | 0.88 | 0.38 |
| σ | RatingScale(Factual): Stimulus (Cold): Prediction(Cold) | -0.1 | 0.036 | -2.8 | 0.0051 |
| ν | Intercept | 1.5 | P.094 | 16 | 1.6e-59 |

| value ~ RateCon * Stimulus * Prediction + Trial Number + re(random = ~ Stimulus \| id), ZOIB(link = logit) | | | | | |
|---|---|---|---|---|---|
| **parameter** | **contrast** | **β** | **SE** | **t** | **p** |
| ν | RatingScale(Factual) | -6.8 | 0.13 | -53 | 0 |
| ν | Stimulus(Cold) | -0.75 | 0.11 | -6.7 | 2.6e-11 |
| ν | Prediction(Cold) | -0.091 | 0.11 | -0.81 | 0.42 |
| ν | Trial Number | -0.00088 | 0.00033 | -2.7 | 0.0076 |
| ν | RatingScale(Factual): Stimulus(Cold) | 1.7 | 0.17 | 9.9 | 3.7e-23 |
| ν | RatingScale(Factual): Prediction(Cold) | 0.76 | 0.17 | 4.5 | 6.9e-06 |
| ν | Stimulus(Cold): Prediction(Cold) | 0.47 | 0.16 | 3 | 0.003 |
| ν | RatingScale(Factual): Stimulus (Cold): Prediction(Cold) | -1.9 | 0.24 | -7.9 | 2.6e-15 |
| τ | Intercept | -6.3 | 0.35 | -18 | 1.8e-71 |
| τ | RatingScale(Factual) | 0.19 | 0.39 | 0.48 | 0.63 |
| τ | Stimulus(Cold) | -1.5 | 0.66 | -2.3 | 0.023 |
| τ | Prediction(Cold) | -0.3 | 0.44 | -0.68 | 0.5 |
| τ | Trial Number | -0.00082 | 0.0011 | -0.73 | 0.46 |
| τ | RatingScale(Factual): Stimulus(Cold) | 2 | 0.74 | 2.7 | 0.006 |
| τ | RatingScale(Factual): Prediction(Cold) | -0.2 | 0.57 | -0.35 | 0.73 |
| τ | Stimulus (Cold): Prediction(Cold) | -1 | 1.3 | -0.83 | 0.41 |
| τ | RatingScale(Factual): Stimulus (Cold): Prediction(Cold) | 1.6 | 1.3 | 1.2 | 0.24 |

**Table 2e,** Accuracy in the following trial given the perceived thermosensory quality of TGI and cue-outcome contingency.

| **predAcc2 ~ coolness * Contingency + Trial Number + random(id), Binomial(link = logit)** | | | | | |
|---|---|---|---|---|---|
| **parameter** | **contrast** | **β** | **SE** | **t** | **p** |
| μ | Intercept | -0.075 | 0.069 | -1.1 | 0.28 |
| μ | Perceived Coldness | 1 | 0.11 | 9.3 | 1.5e-20 |
| μ | Contingency predicts Warm | 0.73 | 0.076 | 9.6 | 1.3e-21 |
| μ | Trial Number | -0.002 | 0.00024 | -8.4 | 7e-17 |
| μ | Perceived Coldness: Contingency predicts Warm | -1.7 | 0.16 | -11 | 1.5e-27 |

**Table 2f,** Response times in the following trial given the perceived thermosensory quality of TGI and cue-outcome contingency.

| predRT2 ~ Perceived Coolness * Contingency + re(random = ~1 \| id), Gamma(link = log) | | | | | |
|---|---|---|---|---|---|
| **parameter** | **contrast** | **β** | **SE** | **t** | **p** |
| μ | Intercept | 0.19 | 0.013 | 15 | 3.3e-48 |
| μ | Perceived Coldness | -0.02 | 0.025 | -0.8 | 0.43 |
| μ | Contingency predicts(Warm) | -0.11 | 0.018 | -5.8 | 5.4e-09 |
| μ | Perceived Coldness: Contingency predicts Warm | 0.14 | 0.037 | 3.7 | 0.00025 |
| σ | Intercept | -0.71 | 0.018 | -40 | 2e-323 |
| σ | Perceived Coldness | -0.04 | 0.035 | -1.2 | 0.25 |
| σ | Contingency predicts Warm | 0.011 | 0.025 | 0.44 | 0.66 |
| σ | Perceived Coldness: Contingency predicts Warm | -0.026 | 0.051 | -0.51 | 0.61 |

**Table 3a,** Effect of prediction uncertainty on accuracy in the current trial.

| predAcc ~ Prediction Uncertainty + Trial Number + random(id), Binomial(link = logit) | | | | | |
|---|---|---|---|---|---|
| **parameter** | **contrast** | **β** | **SE** | **t** | **p** |
| μ | Intercept | 1.5 | 0.027 | 55 | 0 |
| μ | Prediction Uncertainty | -4.9 | 0.13 | -38 | 3.7e-317 |
| μ | Trial Number | -0.00091 | 9e-05 | -10 | 2.7e-24 |

**Table 3b,** Effect of prediction uncertainty on response times in the current trial.

| predRT ~ Prediction Uncertainty + Trial Number + re(random = ~1 \| id), Gamma(link = log) | | | | | |
|---|---|---|---|---|---|
| **parameter** | **contrast** | **β** | **SE** | **t** | **p** |
| μ | Intercept | -0.31 | 0.0069 | -45 | 0 |
| μ | Prediction Uncertainty | 1.8 | 0.033 | 53 | 0 |
| μ | Trial Number | -0.00036 | 2.3e-05 | -16 | 5.8e-55 |
| σ | Intercept | -0.48 | 0.0078 | -61 | 0 |
| σ | Prediction Uncertainty | -0.61 | 0.038 | -16 | 7.5e-60 |
| σ | Trial Number | 0.00028 | 2.7e-05 | 10 | 1.4e-25 |

**Table 4a,** Effects of belief that the next stimulus would be cold (i.e., Belief towards cold) on thermosensory ratings.

| **Rating(ColdBeta \| WarmBeta \| BurningBeta) ~ Belief(Cold) * RatingScale * Stimulus + Trial Number + random(id), ZOIB(link = logit)** | | | | | |
|---|---|---|---|---|---|
| **parameter** | **contrast** | **β** | **SE** | **t** | **p** |
| μ | (Intercept) | -0.36 | 0.014 | -26 | 4.8e-152 |
| μ | Belief To Cold | 0.15 | 0.023 | 6.4 | 1.2e-10 |
| μ | RatingScale(BurnBeta) | -1 | 0.025 | -40 | 0 |
| μ | RatingScale(WarmBeta) | -0.77 | 0.025 | -30 | 6.6e-203 |
| μ | Stimulus(TGI) | -0.32 | 0.022 | -15 | 1.2e-49 |
| μ | Stimulus(Warm) | -0.79 | 0.027 | -30 | 7.1e-192 |
| μ | Trial number | 0.00013 | 3.1e-05 | 4.2 | 2.5e-05 |
| μ | Belief(Cold):RatingScale (BurnBeta) | -0.046 | 0.045 | -1 | 0.31 |
| μ | Belief(Cold):RatingScale (WarmBeta) | -0.19 | 0.045 | -4.1 | 4.1e-05 |
| μ | Belief(Cold):Stim(TGI) | 0.022 | 0.039 | 0.57 | 0.57 |
| μ | Belief(Cold):Stim(Warm ) | -0.26 | 0.046 | -5.7 | 1e-08 |
| μ | RatingScale(BurnBeta): Stim(TGI) | 0.78 | 0.037 | 21 | 1.3e-98 |
| μ | RatingScale(WarmBeta) :Stim(TGI) | 1.4 | 0.034 | 40 | 0 |
| μ | RatingScale(BurnBeta): Stim(Warm) | 0.61 | 0.04 | 15 | 3.7e-52 |
| μ | RatingScale(WarmBeta) :Stim(Warm) | 1.6 | 0.036 | 45 | 0 |
| μ | Belief(Cold):RatingScale (BurnBeta):Stim(TGI) | -0.17 | 0.066 | -2.5 | 0.012 |
| μ | Belief(Cold):RatingScale (WarmBeta):Stim(TGI) | -0.15 | 0.062 | -2.4 | 0.016 |
| μ | Belief(Cold:RatingScale( BurnBeta):Stim(Warm) | 0.12 | 0.071 | 1.6 | 0.1 |

| **Rating(ColdBeta \| WarmBeta \| BurningBeta) ~ Belief(Cold) * RatingScale * Stimulus + Trial Number + random(id), ZOIB(link = logit)** | | | | | |
|---|---|---|---|---|---|
| **parameter** | **contrast** | **β** | **SE** | **t** | **p** |
| μ | Belief(Cold):RatingScale(WarmBeta):Stim(Warm) | 0.22 | 0.064 | 3.4 | 0.00075 |
| σ | (Intercept) | -0.53 | 0.017 | -31 | 1.1e-207 |
| σ | Belief(Cold) | -0.12 | 0.029 | -4.2 | 2.5e-05 |
| σ | RatingScale(BurnBeta) | 0.27 | 0.026 | 10 | 6.7e-25 |
| σ | RatingScale(WarmBeta) | 0.31 | 0.026 | 12 | 3.7e-33 |
| σ | Stim(TGI) | 0.17 | 0.025 | 6.9 | 5.3e-12 |
| σ | Stim(Warm) | 0.46 | 0.026 | 18 | 3.3e-69 |
| σ | Trial Number | -0.00013 | 3.4e-05 | -3.9 | 9.2e-05 |
| σ | Belief(Cold):RatingScale(BurnBeta) | 0.14 | 0.046 | 3.1 | 0.0022 |
| σ | Belief(Cold):RatingScale(WarmBeta) | 0.19 | 0.047 | 4.1 | 4.2e-05 |
| σ | Belief(Cold):Stim(TGI) | 0.12 | 0.045 | 2.6 | 0.01 |
| σ | Belief(Cold):Stimulus(Warm) | 0.082 | 0.046 | 1.8 | 0.079 |
| σ | RatingScale(BurnBeta):Stimulus(TGI) | -0.11 | 0.038 | -2.8 | 0.0051 |
| σ | RatingScale(WarmBeta):Stimulus(TGI) | -0.48 | 0.037 | -13 | 5.1e-37 |
| σ | RatingScale(BurnBeta:Stimulus(Warm) | -0.58 | 0.039 | -15 | 6.9e-51 |
| σ | RatingScale(WarmBeta):Stimulus(Warm) | -0.93 | 0.037 | -25 | 1.7e-141 |
| σ | Belief(Cold):RatingScale(BurnBeta):Stimulus(TGI) | -0.11 | 0.068 | -1.6 | 0.11 |
| σ | Belief(Cold):RatingScale(WarmBeta):Stimulus(TGI) | -0.068 | 0.067 | -1 | 0.31 |

| **Rating(ColdBeta \| WarmBeta \| BurningBeta) ~ Belief(Cold) * RatingScale * Stimulus + Trial Number + random(id), ZOIB(link = logit)** | | | | | |
|---|---|---|---|---|---|
| **parameter** | **contrast** | **β** | **SE** | **t** | **p** |
| σ | Belief(Cold):RatingScale(BurnBeta):Stimulus(Warm) | -0.12 | 0.068 | -1.8 | 0.075 |
| σ | Belief(Cold):RatingScale(WarmBeta):Stimulus(Warm) | -0.14 | 0.065 | -2.2 | 0.027 |
| v | (Intercept) | -4.6 | 0.12 | -38 | 8.9e-322 |
| v | Belief(Cold) | -0.62 | 0.22 | -2.8 | 0.0047 |
| v | RatingScale(BurnBeta) | 4.8 | 0.16 | 30 | 5.9e-203 |
| v | RatingScale(WarmBeta) | 5.5 | 0.16 | 33 | 8.7e-242 |
| v | Stimulus(TGI) | 2.3 | 0.15 | 15 | 3e-53 |
| v | Stimulus(Warm) | 6.3 | 0.17 | 37 | 2.3e-295 |
| v | Trial Number | 0.00097 | 0.00019 | 5 | 5.9e-07 |
| v | Belief(Cold):RatingScale(BurnBeta) | 0.51 | 0.29 | 1.8 | 0.075 |
| v | Belief(Cold):RatingScale(WarmBeta) | 0.95 | 0.31 | 3.1 | 0.002 |
| v | Belief(Cold):Stimulus(TGI) | -0.048 | 0.27 | -0.18 | 0.86 |
| v | Belief(Cold):Stimulus(Warm) | 0.43 | 0.31 | 1.4 | 0.16 |
| v | RatingScale(BurnBeta):Stimulus(TGI) | -4.1 | 0.2 | -20 | 1e-89 |
| v | RatingScale(WarmBeta):Stimulus(TGI) | -7.2 | 0.21 | -34 | 1.2e-246 |
| v | RatingScale(BurnBeta):Stimulus(Warm) | -5.7 | 0.23 | -25 | 5.3e-139 |
| v | RatingScale(WarmBeta):Stimulus(Warm) | -13 | 0.25 | -51 | 0 |
| v | Belief(Cold):RatingScale(BurnBeta):Stimulus(TGI) | 0.36 | 0.37 | 0.96 | 0.34 |

| **Rating(ColdBeta \| WarmBeta \| BurningBeta) ~ Belief(Cold) * RatingScale * Stimulus + Trial Number + random(id), ZOIB(link = logit)** | | | | | |
|---|---|---|---|---|---|
| **parameter** | **contrast** | **β** | **SE** | **t** | **p** |
| ν | Belief(Cold):RatingScale(WarmBeta):Stimulus(TGI) | 0.18 | 0.39 | 0.47 | 0.64 |
| ν | Belief(Cold):RatingScale(BurnBeta):Stimulus(Warm) | -0.035 | 0.41 | -0.085 | 0.93 |
| ν | Belief(Cold):RatingScale(WarmBeta):Stimulus(Warm) | -0.21 | 0.44 | -0.48 | 0.63 |
| τ | (Intercept) | -8.8 | 0.32 | -27 | 9.8e-163 |
| τ | Belief(Cold) | -0.22 | 0.57 | -0.38 | 0.7 |
| τ | RatingScale(BurnBeta) | 0.22 | 0.51 | 0.43 | 0.66 |
| τ | RatingScale(WarmBeta) | -1 | 0.92 | -1.1 | 0.26 |
| τ | Stimulus(TGI) | -1.3 | 0.57 | -2.2 | 0.028 |
| τ | Stimulus(Warm) | -0.95 | 0.58 | -1.6 | 0.1 |
| τ | Trial Number | 0.0018 | 0.00065 | 2.8 | 0.0056 |
| τ | Belief(Cold):RatingScale(BurnBeta) | -1.9 | 1.1 | -1.8 | 0.071 |
| τ | Belief(Cold):RatingScale(WarmBeta) | -3.6 | 2.4 | -1.5 | 0.14 |
| τ | Belief(Cold):Stimulus(TGI) | 1.1 | 1 | 1.1 | 0.26 |
| τ | Belief(Cold):Stimulus(Warm) | 0.25 | 1 | 0.24 | 0.81 |
| τ | RatingScale(BurnBeta):Stimulus(TGI) | 1.8 | 0.76 | 2.4 | 0.017 |
| τ | RatingScale(WarmBeta):Stimulus(TGI) | 2.7 | 1.1 | 2.5 | 0.014 |
| τ | RatingScale(BurnBeta):Stimulus(Warm) | -0.72 | 0.87 | -0.83 | 0.41 |
| τ | RatingScale(WarmBeta):Stimulus(Warm) | 1.2 | 1.1 | 1.1 | 0.29 |

| Rating(ColdBeta \| WarmBeta \| BurningBeta) ~ Belief(Cold) * RatingScale * Stimulus + Trial Number + random(id), ZOIB(link = logit) | | | | | |
|---|---|---|---|---|---|
| **parameter** | **contrast** | **β** | **SE** | **t** | **p** |
| τ | Belief(Cold):RatingScale (BurnBeta):Stimulus(TGI) | 1.1 | 1.5 | 0.75 | 0.45 |
| τ | Belief(Cold):RatingScale (WarmBeta):Stimulus(TGI) | 2.6 | 2.6 | 1 | 0.31 |
| τ | Belief(Cold):RatingScale (BurnBeta):Stimulus(Warm) | 3.1 | 1.6 | 1.9 | 0.052 |
| τ | Belief(Cold):RatingScale (WarmBeta):Stimulus(Warm) | 3.4 | 2.7 | 1.3 | 0.21 |

**Table 4b,** Effects of estimation uncertainty on burning ratings.

| BurningBeta ~ Estimation Uncertainty * Stimulus + Trial Number + re(random = ~Stimulus:sa2 \| id), ZOIB(link = logit) | | | | | |
|---|---|---|---|---|---|
| **parameter** | **contrast** | **β** | **SE** | **t** | **p** |
| μ | Intercept | -1 | 0.017 | -62 | 0 |
| μ | Estimation Uncertainty | 0.013 | 0.006 | 2.1 | 0.032 |
| μ | Stimulus(Cold) | -0.34 | 0.021 | -16 | 9.5e-61 |
| μ | Stimulus(Warm) | -0.6 | 0.021 | -28 | 3.6e-175 |
| μ | Trial Number | 0.00071 | 5.8e-05 | 12 | 8.9e-35 |
| μ | Estimation Uncertainty: Stimulus(Warm) | -0.065 | 0.0097 | -6.7 | 2.1e-11 |
| μ | Estimation Uncertainty: Stimulus(Cold) | -0.034 | 0.0098 | -3.5 | 0.0005 |
| σ | Intercept | -0.98 | 0.012 | -82 | 0 |
| σ | Estimation Uncertainty | -0.026 | 0.005 | -5.2 | 1.6e-07 |
| σ | Stimulus(Cold) | 0.04 | 0.015 | 2.7 | 0.006 |
| σ | Stimulus(Warm) | -0.0021 | 0.015 | -0.14 | 0.89 |

| BurningBeta ~ Estimation Uncertainty * Stimulus + Trial Number + re(random = ~Stimulus:sa2 \| id), ZOIB(link = logit) | | | | | |
|---|---|---|---|---|---|
| **parameter** | **contrast** | **β** | **SE** | **t** | **p** |
| σ | Trial Number | -6.2e-05 | 3.8e-05 | -1.6 | 0.11 |
| σ | Estimation Uncertainty: Stimulus(Warm) | -0.019 | 0.0074 | -2.6 | 0.011 |
| σ | Estimation Uncertainty: Stimulus(Cold) | -0.013 | 0.0074 | -1.8 | 0.069 |
| ν | Intercept | -0.37 | 0.12 | -3.2 | 0.0016 |
| ν | Estimation Uncertainty | -0.22 | 0.049 | -4.5 | 8.1e-06 |
| ν | Stimulus(Cold) | 1.8 | 0.14 | 13 | 7.1e-38 |
| ν | Stimulus(Warm) | 3.2 | 0.14 | 23 | 2.1e-120 |
| ν | Trial Number | -0.00044 | 0.00034 | -1.3 | 0.2 |
| ν | Estimation Uncertainty: Stimulus(Warm) | 0.28 | 0.067 | 4.2 | 3.1e-05 |
| ν | Estimation Uncertainty: Stimulus(Cold) | 0.21 | 0.067 | 3.2 | 0.0015 |
| τ | Intercept | -6.5 | 0.31 | -21 | 1.4e-94 |
| τ | Estimation Uncertainty | 0.26 | 0.1 | 2.5 | 0.011 |
| τ | Stimulus(Cold) | -0.49 | 0.44 | -1.1 | 0.26 |
| τ | Stimulus(Warm) | -0.46 | 0.44 | -1.1 | 0.29 |
| τ | Trial Number | 0.0061 | 0.0012 | 5.1 | 3.7e-07 |
| τ | Estimation Uncertainty: Stimulus(Warm) | -0.86 | 0.32 | -2.7 | 0.0071 |
| τ | Estimation Uncertainty: Stimulus(Cold) | -0.9 | 0.32 | -2.8 | 0.0054 |

### *Supplementary Figures*

To ensure the robustness of our models, we conducted parameter recovery analysis. This analysis revealed that the 3-level Hierarchical Gaussian Filter model could not adequately recover the two parameters (i.e., θ and κ) governing the third level. Consequently, this model was not included in neither model comparison nor model selection. More details are reported in Supplementary Figure 4.

**Supplementary Figure 1: Parameter recovery analysis of the 2-level Hierarchical Gaussian Filter learning model.**

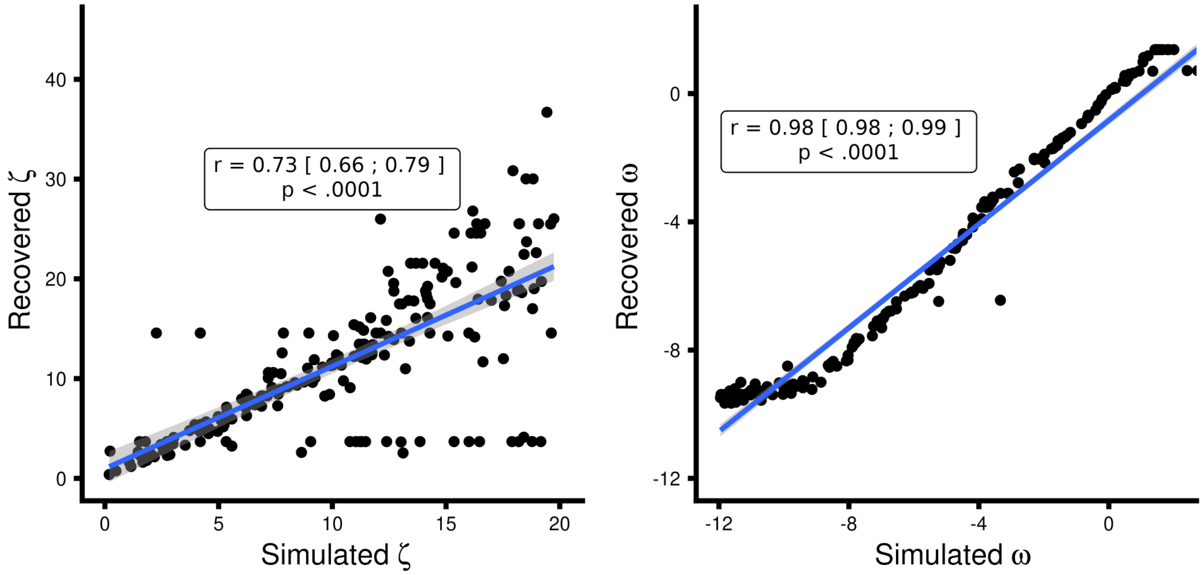


X-axis presenting the simulated values and the y-axis being the estimated / recovered value. Very wide non-informative priors were set for both parameters i.e., $\omega \sim N\left(e^{-3}, 16\right)$ and $\zeta \sim N(log(5), 3)$. Note we display $N\left(\mu, \sigma^2\right)$ as the HGF toolbox. All parameters were simulated from a uniform distribution in the range seen in the plot below.

**Supplementary Figure 2: Parameter recovery analysis of the Rescorla-Wagner learning model.**

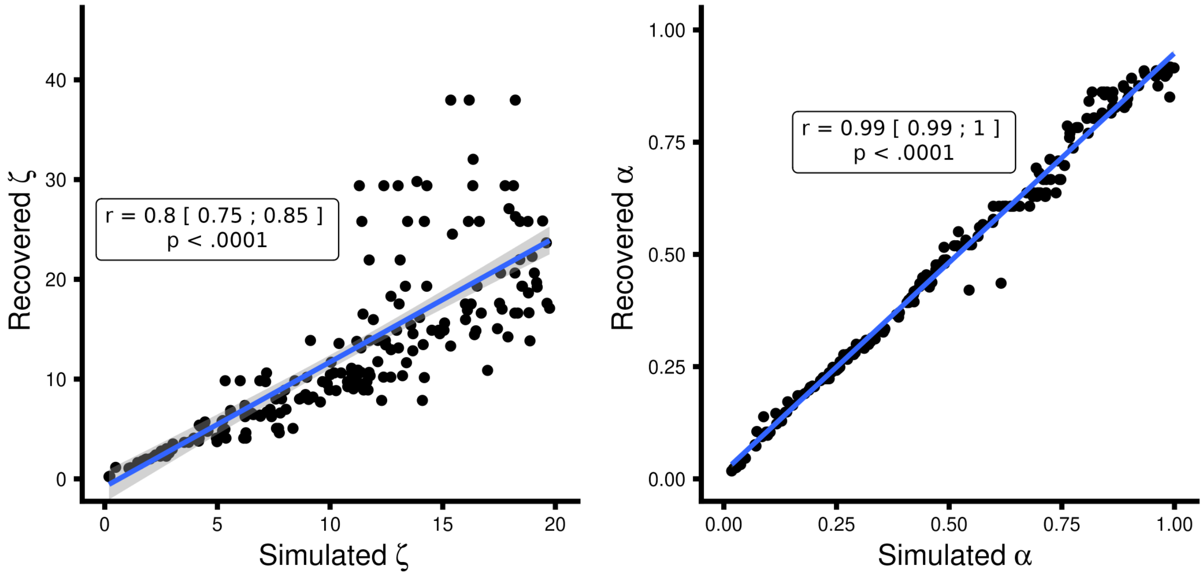


X-axis presenting the simulated values and the y-axis being the estimated / recovered value. Very wide non-informative priors were set for both parameters i.e., $\alpha \sim N(S(0.5), 2)$ and $\zeta \sim N(log(5), 3)$, note we display $N(\mu, \sigma^2)$ as the HGF toolbox. Where $S(x)$ represents the sigmoid transformation i.e. $S(x) = \frac{1}{1+exp^{-x}}$ . All parameters were simulated from a uniform distribution in the range seen in the plot below

**Supplementary Figure 3: Parameter recovery analysis of the Sutton K1 learning model.**

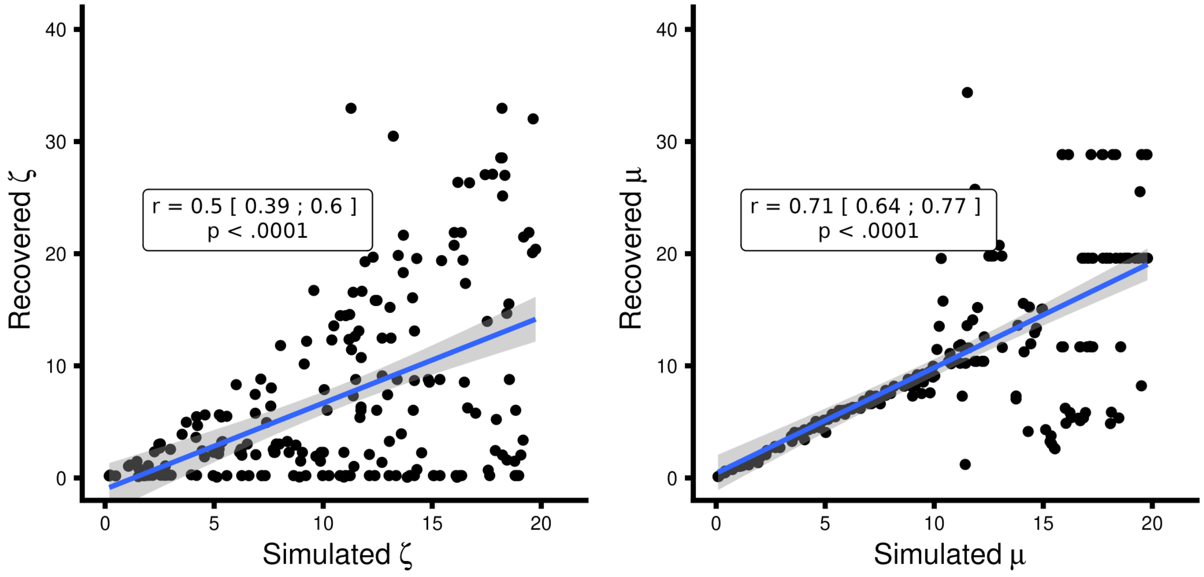


X-axis presenting the simulated values and the y-axis being the estimated / recovered value. Very wide non-informative priors were set for both parameters i.e., $\mu \sim N(log(3), 16)$ and $\zeta \sim N(log(5), 3)$, Note we display $N\left(\mu, \sigma^2\right)$. Where $S(x)$ represents the sigmoid transformation i.e. $S(x) = \frac{1}{1+exp^{-x}}$. All parameters were simulated from a uniform distribution in the range seen in the plot below

**Supplementary Figure 4: Parameter recovery analysis of the 3-level Hierarchical Gaussian Filter learning model.**

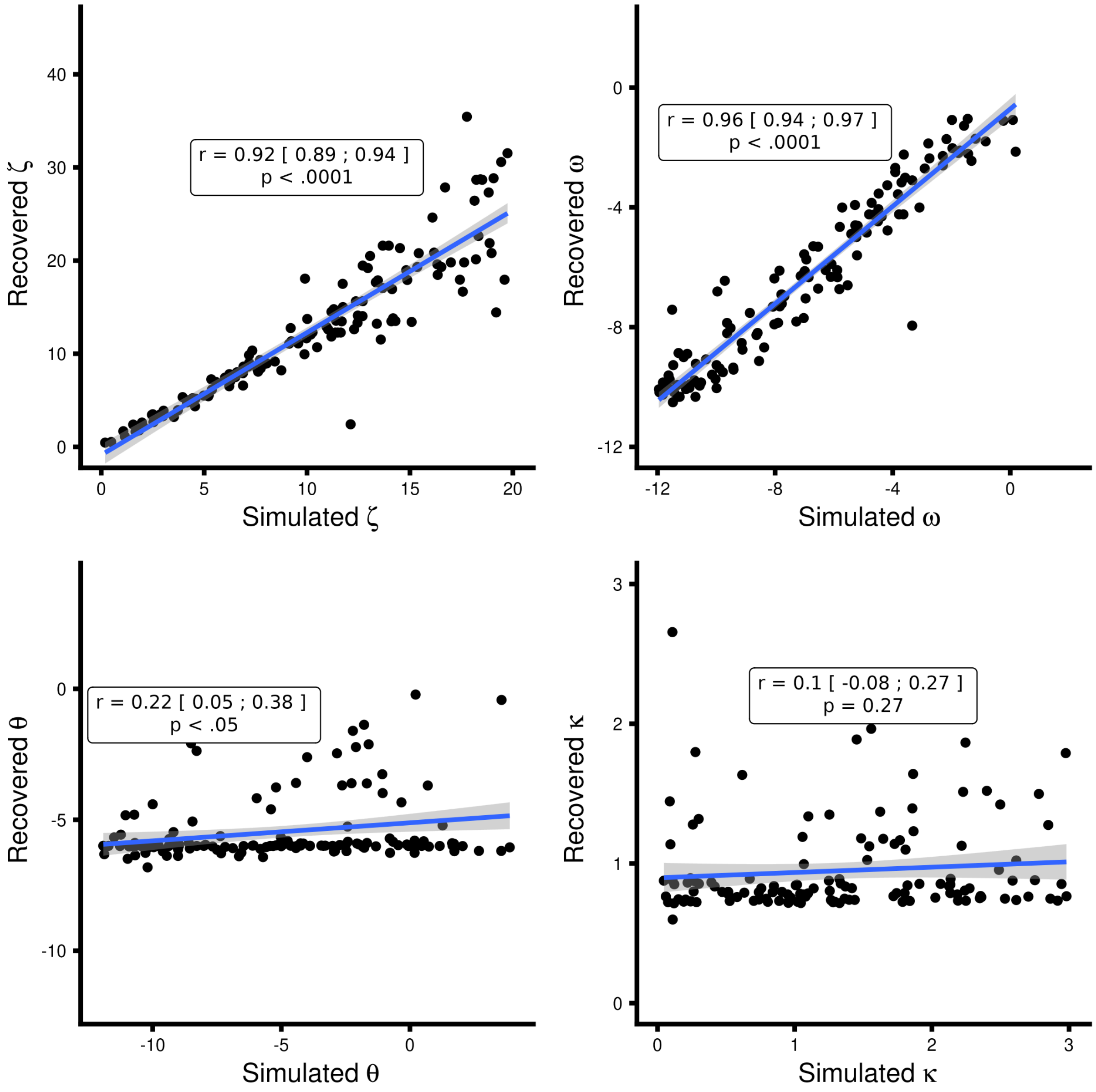


X-axis presenting the simulated values and the y-axis being the estimated / recovered value. Very wide non-informative priors were set for both parameters i.e., $\omega \sim N\left(e^{-3}, 16\right)$ and $\zeta \sim N(log(5), 3)$, $\theta \sim N\left(e^{-6}, 16\right)$ and $\kappa \sim N\left(e^{log(1)}, 1\right)$. Note we display $N\left(\mu, \sigma^2\right)$ as the HGF toolbox. All parameters were simulated from a uniform distribution in the range seen in the plot below. Due to the very poor recovery of the third level parameters i.e. κ and θ the 3-level HGF model was not used in model comparison.

The parameter recovery analysis demonstrated that the 2-level HGF, the Rescorla Wagner and the Sutton k1 learning models successfully recovered their respective parameters with acceptable precision. However, the 3-level HGF failed to recover any parameters at its third level, making it unsuitable for further analysis in this context. The outcomes of the parameter recovery were then utilized to establish suitable priors for subsequent model recovery analyses. For further details, including comprehensive plots that illustrate the evaluation of the priors used in our simulations, readers are directed to the Matlab directory in the GitHub repository linked to this study.

**Supplementary Figure 4: Model recovery analyses.**

| | Simulated | | |
|---|---|---|---|
| | HGF | RW | Sutton |
| Recovered HGF | 184 | 36 | 43 |
| Recovered RW | 11 | 150 | 1 |
| Recovered Sutton | 5 | 14 | 156 |

**Supplementary Figure 5: Model selection analysis using random effects on log model evidence.**

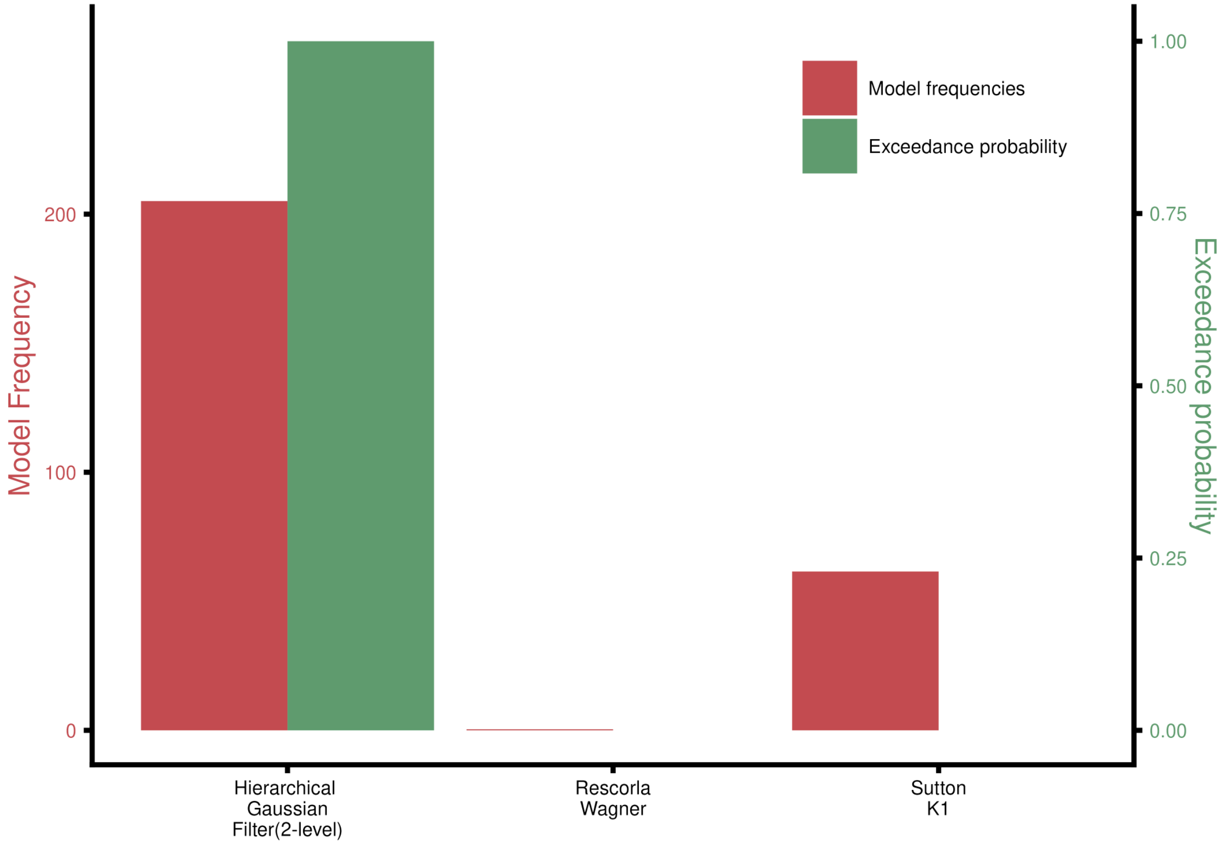


The Hierarchical Gaussian Filter outperformed the fixed learning rate model, Rescorla–Wagner, and the variable-learning-rate non-Bayesian model Sutton K1.
